# Genetic Inactivation of β-Catenin Attenuates and Its Activation Aggravates Desmoplakin Cardiomyopathy

**DOI:** 10.1101/2023.03.02.530831

**Authors:** Melis Olcum, Siyang Fan, Leila Rouhi, Sirisha Cheedipudi, Benjamin Cathcart, Hyun-Hwan Jeong, Zhongming Zhao, Priyatansh Gurha, Ali J. Marian

## Abstract

**Aim:** Mutations in the *DSP* gene encoding desmoplakin, a constituent of the desmosomes at the intercalated discs (IDs), cause a phenotype that spans arrhythmogenic cardiomyopathy (ACM) and dilated cardiomyopathy (DCM). It is typically characterized by biventricular enlargement and dysfunction, severe myocardial fibrosis, cell death, and arrhythmias.

The canonical WNT (cWNT)/β-catenin signaling pathway is implicated in the pathogenesis of ACM. Given that β-catenin, an indispensable co-transcriptional regulator of the cWNT pathway, is also a member of the IDs, we genetically inactivated or activated β-catenin to determine its role in the pathogenesis of the desmoplakin cardiomyopathy.

**Methods and Results:** The *Dsp* gene was conditionally deleted in cardiac myocytes concomitant with the genetic inactivation or activation of β-catenin using the tamoxifen-inducible MerCreMer mice. Inactivation and activation of β-catenin were achieved upon deletion of its transcriptional domain and degrons, respectively. Analysis of cardiac myocytes transcripts and proteins showed marked dysregulation of the cWNT/β-catenin pathway in the DSP-deficient mouse cardiac myocytes (*Myh6-Mcm*^Tam^:*Dsp*^F/F^), as indicated by increased expression of cWNT/β-catenin targets along with its inhibitors and isoforms of its key co-effectors. Genetic inactivation of β-catenin in the *Myh6-Mcm*^Tam^:*Dsp*^F/F^ mice prolonged survival, improved cardiac function, reduced cardiac arrhythmias, and attenuated myocardial fibrosis, and cell death caused by apoptosis, necroptosis, pyroptosis, i.e., PANoptosis, whereas its activation had the opposite effects. Inactivation of β-catenin was associated with partial restoration of the suppressed genes involved in OXPHOS, whereas its activation has the opposite effect. The beneficial effects were independent of the changes in the transcript levels of the cWNT target genes.

**Conclusion:** The cWNT/β-catenin was markedly dysregulated in the cardiac myocytes from a mouse model of DC. Inactivation of β-catenin attenuated the phenotype partly through the recovery of OXPHOS genes whereas its activation had deleterious effects. The findings suggest suppression of β-catenin might be beneficial in desmoplakin-cardiomyopathy.

**Summary:** Genetic inactivation of β-catenin improved desmoplakin cardiomyopathy, in part through the restoration of expression of genes involved in oxidative phosphorylation, whereas its activation was deleterious.

## INTRODUCTION

Hereditary cardiomyopathies comprise a genetically and phenotypically heterogenous group of myocardial diseases wherein the primary defect is in cardiac myocytes, the ensuing phenotype, however, is multicellular involving cellular constituents of the myocardium. (1-4) Despite their heterogeneity, hereditary cardiomyopathies exhibit partial genetic and phenotypic overlaps. Among the three most common forms of hereditary cardiomyopathies, namely hypertrophic cardiomyopathy (HCM), dilated cardiomyopathy (DCM), and arrhythmogenic cardiomyopathy (ACM), the latter two exhibit considerable overlaps and are at times indistinguishable. Consequently, terms such as arrhythmogenic left dominant cardiomyopathy and arrhythmogenic DCM are used to describe overlapping phenotypes. (5, 6)

DCM is characterized by cardiac dilatation and reduced LVEF and is caused by mutations in genes coding for the cytoskeletal or sarcomere proteins. (1) ACM, in its classic form, predominantly involves the right ventricle and is referred to as arrhythmogenic right ventricular cardiomyopathy (ARVC) and is primarily a disease of the intercalated disk (ID) proteins, mostly desmosome proteins. (3) Among the genes known to cause ACM, the *DSP* gene, which encodes desmoplakin (DSP), causes a somewhat distinct phenotype that straddles between DCM and ACM (5, 7) The phenotypic overlap likely reflects the topographic location of DSP in the intracellular component of the desmosome structures where it interacts with the cytoskeletal proteins, such as cardiac α-actin, which are known to cause DCM. (8, 9) The phenotype caused by the *DSP* mutations is somewhat distinct, as it typically involves both ventricles, leading to biventricular dilatation and dysfunction, ventricular tachycardia, severe myocardial fibrosis, and apoptosis. (6, 7, 10, 11) Consequently, the phenotype is recognized as desmoplakin cardiomyopathy. (6, 7, 10, 11)

The cardiac IDs are comprised of multi-functional proteins that attach the adjacent cardiac myocytes and provide mechanical integrity to the myocardium, enable rapid electric conduction throughout the myocardium, and serve as a signaling hub responding to external or internal mechanical stress. (12-16) The β-catenin (CTNNB1), which is the indispensable transcriptional co-effectors of the cWNT pathway is an abundant constituent of the adherens junction at the IDs, where it interacts with the mechano-sensing proteins N-cadherin and α-catenin. (17, 18) Given the intricate interplays between the β-catenin and protein constituents of desmosomes, the cWNT/β-catenin is implicated in the pathogenesis of ACM and has emerged as a target of experimental therapy. (16, 19-24) The specific role of β-catenin, however, in the pathogenesis of ACM has remained unexplored. Therefore, we utilized genetic loss- and gain-of-function (LoF and GoF, respectively) approaches to inactivate or activate β-catenin in cardiac myocytes in a mouse model of DC caused by cardiac myocyte-specific DSP deficiency and determine the phenotypic effects.

## METHODS

### Data sharing

RNA-Seq data have been submitted to GEO (GSE180972).

### Regulatory approvals

The use of mice in these studies was approved by the Institutional Animal Care and Use Committee (AWC-21-0015) and was per the NIH Guide for the Care and Use of Laboratory Animals.

### Anesthesia and euthanasia

Isoflurane was used to induce (3%) and maintain (0.5%) anesthesia in mice. Euthanasia was accomplished after placing the mice under 100% CO2 inhalation and cervical dislocation.

### The *Myh6-Mcm*:*Dsp*^F/F^ mice

The *Dsp* gene was specifically deleted in the post-natal cardiac myocytes upon daily intra-peritoneal injection of 5 consecutive doses of tamoxifen (30 mg/Kg/d) to the *Myh6-Mcm*:*Dsp*^F/F^ mice starting at post-natal day 14, as published. (11)

### Genetic activation or inactivation of β-catenin (CTNNB1) in cardiac myocytes in the post-natal mice

The mouse models of tamoxifen-inducible Cre-mediated β-catenin activation and inactivation have been published. (25) In brief, to activate β-catenin, the floxed exon 3 of the *Ctnnb1* gene, which encodes the GSK3β phosphorylation sites (Thr41, Ser37, and Ser33), necessary for its degradation by ubiquitinoylation (degron), was deleted. (25-27) The deletion leads to the expression of a stable and ∼ 10 kDa (80 amino acids) smaller β-catenin protein. The mouse represents the GoF of β-catenin (*Ctnnb1*^GoF^). To inactivate β-catenin, floxed exons 8 to 13 of the *Ctnnb1* gene, which encodes the trans-activator motif comprised of amino acids 361-692, were deleted. (25, 28) The deletion leads to the expression of truncated mRNA and protein, which are unstable and degraded. (28) The mouse represents the LoF of β-catenin (*Ctnnb1*^LoF^).

### Inducible activation or inactivation of β-catenin concomitant with deletion of the *Dsp* gene in cardiac myocytes in the post-natal mice

To delete *Dsp* and concomitantly activate or inactivate β-catenin in cardiac myocyte, *Myh6-Mcm*^Tam^:*Dsp*^F/F^:*Ctnnb1*^LoF^ and *Myh6-Mcm*^Tam^:*Dsp*^F/F^:*Ctnnb1*^GoF^ mice were generated and injected daily with tamoxifen (30 mg/kg/d) starting at P14 for 5 consecutive days (Online Figure 1). The P14 time point was chosen to avoid the confounding effects of possible activation or suppression of the cWNT/ β-catenin pathway in the first 2 post-natal weeks when the proliferative capacity of myocytes progressively ceases. (29, 30) The approach also avoids embryonic lethality, which occurs upon homozygous deletion of the *Dsp* gene. (16, 31)

**Figure 1.**
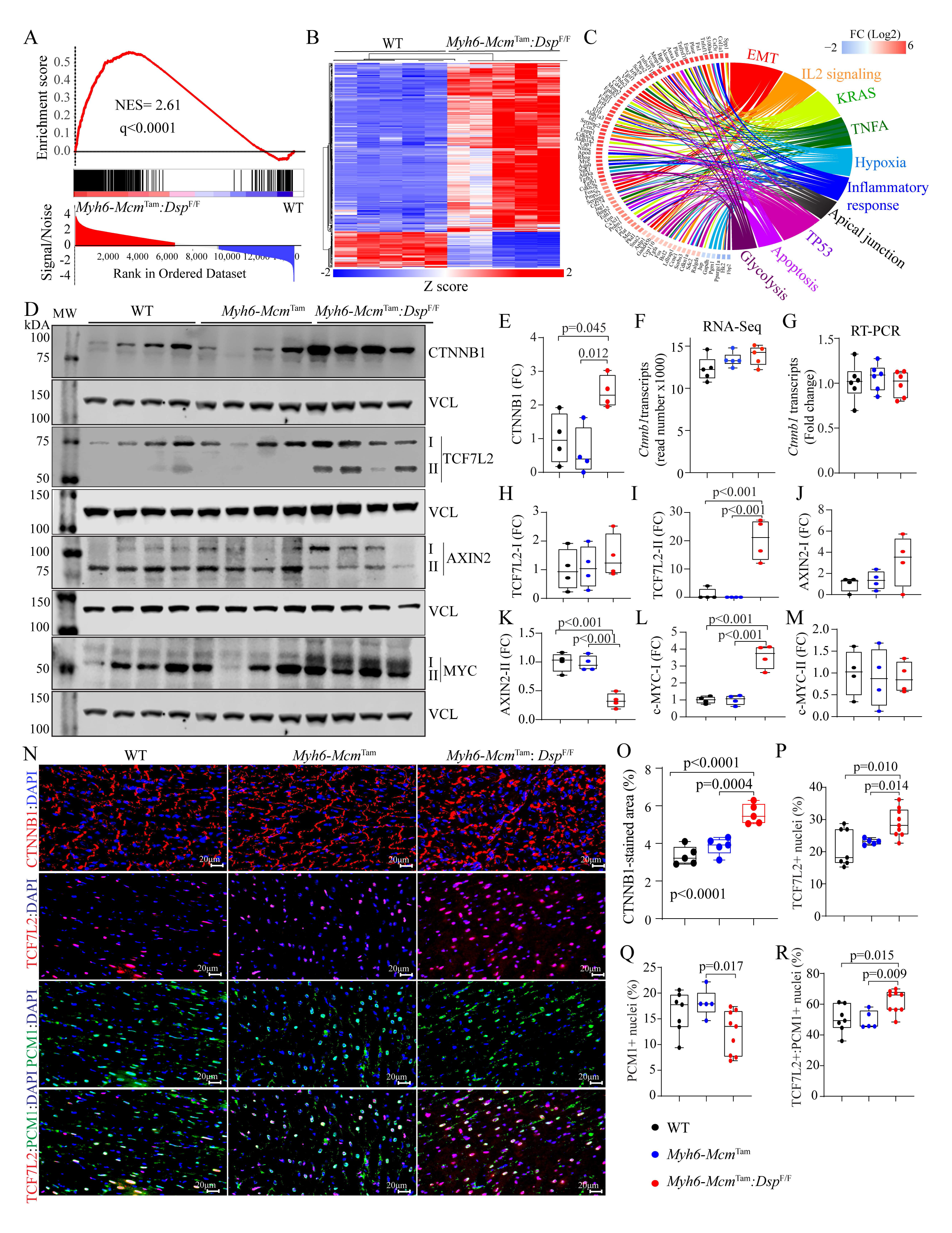
Dysregulated canonical WNT (cWNT) pathway in cardiac myocytes in a mouse model of desmoplakin cardiomyopathy. **A.** Gene set enrichment analysis (GSEA) predicting activation of the cWNT/TCF7L2/β-catenin target genes in the *Myh6-Mcm*^Tam^*:Dsp*^F/F^ mouse cardiac myocytes. **B.** Heat map of the cWNT target genes. **C.** Circos map showing pathways predicted to be affected by the cWNT target genes in the *Myh6-Mcm*^Tam^*:Dsp*^F/F^ mouse cardiac myocytes. **D.** Immunoblots showing protein levels of selected key molecules in the cWNT pathway, showing either upregulation of the main or the alternative isoforms. **E.** Quantification of the β-catenin protein levels. **F** and **G**. Transcript levels of the *Ctnnb1* gene in the RNA-Seq data and determined by RT-PCR in an independent set of RNA extracts. **H-M**. Quantitative data corresponding to TCF7L2 and MYC proteins and their isoforms, which were shown in panel D. **N-R**. Immunofluorescence panels showing expression of β-catenin (CTNNB1), TCF7L2, and PCM1 in thin myocardial sections along with the corresponding quantitative data.

### Genotypes

The mice were genotyped by PCR using genomic DNA isolated from the tail, as described. (16, 32) The list of oligonucleotide primers used for the genotyping is provided in Online Table 1.

**Table 1.** Effects of Genetic Inactivation or Activation of β-Catenin on Echocardiographic Indices of Cardiac Size and Function

|  | WT | *Myh6-Mcm*<sup>Tam</sup>:*Dsp*<sup>F/F</sup> | *Myh6-Mcm*<sup>Tam</sup>:*Dsp*<sup>F/F</sup>:*Ctnnb1*<sup>LoF</sup> | *Myh6-Mcm*<sup>Tam</sup>:*Dsp*<sup>F/F</sup>:*Ctnnb1*<sup>GoF</sup> | p |
| --- | --- | --- | --- | --- | --- |
| N | 18 | 18 | 14 | 14 | NA |
| M/F | 9/9 | 9/9 | 7/7 | 7/7 | >0.9999* |
| Age (days) | 28.28±0.46 | 28.06±0.24 | 28.43±0.65 | 28.50±0.85 | 0.1814# |
| Body weight (g) | 17.71±2.12 | 13.72±2.38 | 13.47±1.99 | 12.00±1.83 | <0.0001 |
| HR (bpm) | 657.71±31.20 | 617.54±52.65 | 654.52±45.42 | 603.72±49.84 | 0.022 |
| IVST (mm) | 0.37±0.03 | 0.29±0.03 | 0.30±0.03 | 0.24±0.02 | <0.0001 |
| LVPWT (mm) | 0.36±0.03 | 0.29±0.03 | 0.30±0.03 | 0.25±0.02 | <0.0001 |
| LVEDD (mm) | 2.91±0.24 | 3.80±0.45 | 3.47±0.24 | 4.15±0.58 | <0.0001<sup>#</sup> |
| LVEDDI (mm/g) | 0.17±0.02 | 0.29±0.07 | 0.26±0.04 | 0.36±0.10 | <0.0001<sup>#</sup> |
| LVESD (mm) | 1.57±0.14 | 3.06±0.58 | 2.60±0.29 | 3.62±0.59 | <0.0001<sup>#</sup> |
| LVESDI (mm/mg) | 0.09±0.01 | 0.23±0.07 | 0.20±0.04 | 0.31±0.09 | <0.0001<sup>#</sup> |
| FS (%) | 45.94±2.01 | 19.94±8.07 | 25.01±6.03 | 13.12±3.07 | <0.0001<sup>#</sup> |
| LV Mass (mg) | 19.74±3.99 | 24.47±5.35 | 21.61±2.85 | 24.21±7.19 | 0.016<sup>#</sup> |
| LVMI (mg/g) | 1.11±0.16 | 1.84±0.55 | 1.63±0.30 | 2.12±0.96 | <0.0001<sup>#</sup> |

**Abbreviations:** WT: Wild type; *Myh6-Mcm*: Myosin heavy chain 6; Mer Cre Mer, *Dsp*: Desmoplakin gene; *Ctnnb1*: β-catenin gene, LOF: Loss of function β-catenin, GOF: Gain of function β-catenin, F/M: Female/Male; g; grams; HR, heart rate; bpm, beats per minute; IVST Diastolic interventricular septum thickness; LVPWT, posterior wall thickness during diastole; LVEDD, left ventricular end diastolic diameter; LVEDDi, LVEDD divided by the body weight; LVESD, left ventricular end systolic diameter; LVESDI, LVESD divided by body weight; FS, Left ventricular fractional shortening; LVM, left ventricular mass; LVMI, LVM divided by the body weight.
**Symbols:** \* Chi square p value, # p value derived by Kruskal-Wallis test.

### Survival analysis

Kaplan Meier survival plots were constructed in the main study groups, namely the wild type (WT), *Myh6-Mcm*^Tam^:*Dsp*^F/F^, *Myh6-Mcm*^Tam^:*Dsp*^F/F^:*Ctnnb1*^LoF,^ and *Myh6-Mcm*^Tam^:*Dsp*^F/F^:*Ctnnb1*^GoF^ using GraphPad Prism 9 software (https://www.graphpad.com/). The survival data in *Myh6-Mcm*^Tam^, *Myh6-Mcm*^Tam^:*Ctnnb1*^LoF,^ and *Myh6-Mcm*^Tam^:*Ctnnb1*^GoF^ have been published and were not included to avoid redundancy. (25, 33)

### Gross morphology

Body and heart weight were measured at 4 weeks of age, which is the time point that most phenotypes were analyzed. The heart weight was indexed to body weight and compared among the groups.

### Echocardiography

Echocardiography was performed on 4-week-old mice using a Vevo 1100 ultrasound imaging system (FUJIFILM VisualSonics Inc., Toronto, ON, Canada), as published. (25, 33, 34) In brief, cardiac images were obtained in the M mode using a B-mode guide at the level of left ventricular (LV) papillary muscles in mice anesthetized with 1% isoflurane. LV anterior wall thickness (AWT), posterior wall thickness (PWT), end-systolic diameter (ESD), and end-diastolic diameter (EDD) were measured using the leading-edge method. The LV fractional shortening (LVFS) and mass were calculated from the m-mode images, as published. (25, 33, 34)

### Rhythm monitoring

Two surface leads were placed over the mouse’s chest and the cardiac rhythm was monitored for about an hour in each mouse under 1% isoflurane anesthesia, as published. (11) (35) A Power Lab 4/30 data acquisition system was used and the data were analyzed using Lab Chart7 software (ADInstruments, Colorado Springs, CO, USA).

### Cardiac myocyte isolation

Cardiac myocytes were isolated from 4-week-old mice as published. (11, 36) In brief, mice were anesthetized, and the heart was excised, mounted onto a Langendorff perfusion system, and perfused with a type 2 collagenase digestion buffer (2.4 mg/ml concentration) at a flow rate of 4 ml/min (Worthington Cat# LS004176). The large blood vessels and the atria were excised from the ventricles and minced in a stop buffer containing 10% calf serum, 12.5μM CaCl_2_, and 2 mM ATP. This was followed by filtration of the cell suspension through a 100 µm cell strainer and centrifugation at 20g to precipitate cardiac myocytes. The isolated cells were treated with CaCL_2_ added to the stop buffer in a stepwise increase from a molar concentration of 100 μM to 900 μM. The isolated myocytes were suspended either in a Qiazol reagent (Qiagen Cat# 79306) for RNA extraction or in a protein extraction buffer for immunoblotting.

### Immunoblotting

Myocardial and cardiac myocyte proteins were analyzed for the expression of selected proteins by immunoblotting. In brief, ventricular tissues or cardiac myocytes were homogenized and dissolved in a RIPA buffer (Cat# 974821) containing 0.5% SDS. (37) The buffers contained protease and phosphatase inhibitors (Roche, Cat#04693159001 and Cat#04906837001, respectively). The lysates were sonicated briefly using a Bioruptor Pico (Diagenode) and centrifuged at 13,000 rpm to precipitate the protein. Aliquots of 30 to 50μg protein extracts were loaded onto an SDS polyacrylamide gel, electrophoresed, and transferred onto nitrocellulose membranes. The membranes were probed with antibodies against the target proteins followed by incubation with the corresponding secondary antibodies. The signals were detected using the ECL western blotting detection kit (Amersham Cat# RPN2106) and the images were collected using the LiCOR (Odyssey) imaging system.

### Reverse transcription-polymerase chain reaction (RT-PCR)

Transcript levels of selected genes were quantified by RT-PCR. In brief, total cardiac myocyte RNA was extracted using a miRNeasy Mini Kit (Qiagen, cat # 217006) and treated with DNase I (Qiagen, cat# 79254). Approximately 1μg of total RNA was used for reverse transcription, which was performed using random primers and a high-capacity cDNA synthesis kit (Applied Biosystems cat# 4368814). The SYBR green probes or TaqMan assays were used in duplicate to determine the transcript levels of the selected genes, which were normalized to the *Gapdh* transcript levels. Changes in the transcript levels were calculated using the ΔΔCT method and presented as fold changes relative to the values in the WT mice. The primers used in the study are listed in the Online table 1.

### RNA-Sequencing (RNA-Seq)

RNA-Seq was performed on ribosome-depleted ventricular cardiac myocyte RNA extracts, as published. (11, 24, 25, 33) In brief, total RNA was extracted using the miRNeasy Mini Kit (Qiagen, Cat # 217004), and extracts with an RNA Integrity Number (RIN) value of >8, determined using an Agilent Bioanalyzer RNA chip, were used to construct sequencing libraries. Strand-specific sequencing libraries were generated using TruSeq stranded total RNA library preparation kit (Illumina Inc. Cat # 20020596) and sequenced as 75bp paired-end reads on an Illumina instrument.

### Myocardial fibrosis

Myocardial fibrosis was assessed by multiple methods, including calculation of collagen volume fraction (CVF) from thin myocardial sections stained with picrosirius red, quantification of the transcript of genes involved in fibrosis by RT-PCR and selected proteins by western blotting and immunofluorescence staining, as published. (11, 25, 33, 38) Likewise, fibrosis markers were analyzed in the cardiac myocyte RNA-Seq dataset.

### Myocyte cross-sectional area (CSA)

To identify cardiac myocytes, thin myocardial sections were stained with an antibody against pericentriolar material protein 1 (PCM1), which mainly marks the myocyte nuclei in the heart (Sigma Cat# HPA023370), as published. (25, 32, 35, 39) The sections were co-stained with wheat germ agglutinin (WGA) conjugated to Texas red (concentration 1μg/ml, Thermo Fisher Scientific, Cat#W21405)) and DAPI to mark the interstitium and nuclei, respectively. The number of myocytes was calculated along with the areas not stained with WGA to determine the myocyte CSA.

### TUNEL assay

The TUNEL assay was performed to detect apoptosis using the terminal deoxynucleotidyl transferase dUTP nick end labeling (TUNEL) assay and In-Situ cell death detection Fluorescein kit (Roche Cat#11684795910), as published. (34, 36, 37)

### Bioinformatics and statistics

The RNA-Seq reads were aligned to the mouse reference genome build mm10 using STAR. (40) The uniquely aligned read pairs were annotated using GENCODE gene model (https://www.gencodegenes.org/mouse/). Read counts were determined using the featureCount tool and normalized using the Remove Unwanted Variation (RUVr) method. (41) The differentially expressed genes (DEGs) were identified using the DESeq2 program. (42) Benjamini-Hochberg method was used to calculate the false discovery rate (FDR) and a cut-off level of <0.05 were considered significant. Normalized count per million (CPM) values were used to generate the heatmaps and volcano plots using the Rstudio (www.rstudio.com) package. The Circos maps were plotted using the GOCHORD function in the R package.

Gene Set Enrichment Analysis (GSEA, version 2.2.3, http://software.broadinstitute.org/gsea/) was utilized to identify the dysregulated biological pathways. Normalized Enrichment Score (NES), determined from the Molecular signature database (MSigDB) 3.0, was used to curate gene sets to identify the Hallmark canonical pathways. The DEGs were also analyzed using the Upstream regulator analysis function of the Ingenuity Pathway Analysis software (IPA®, QIAGEN Redwood City) to predict the altered transcriptional regulators. The targets of the specific transcriptional regulators were identified using the IPA program and those with a p-value of <0.05 for overlap with the IPA target genes and a predicted Z score of < -2 or > 2 were considered dysregulated.

The conventional statistical analysis was performed as published. (11, 43) The Gaussian distribution of each set of data was analyzed by the Shapiro-Wilk normality test and the normally distributed were compared between two groups by the t-test and among multiple groups by one-way ANOVA followed by Bonferroni pairwise comparisons. Data that deviated from a Gaussian distribution were compared by the Kruskal-Wallis test. The categorical data were compared by the Fisher exact or the Chi-Square test. Kaplan-Meier survival plots were compared using the Log-rank test. Statistical analyses were performed using Graph pad Prism 9 or STAT IC, 15.1.

## RESULTS

### Study groups

Seven genotypes were generated, which were comprised of wild type (WT), *Myh6-Mcm*^Tam^, *Myh6-Mcm*^Tam^:*Dsp*^F/F^, *Myh6-Mcm*^Tam^:*Ctnnb1*^LoF^, *Myh6-Mcm*^Tam^:*Ctnnb1*^GoF^, *Myh6-Mcm*^Tam^:*Dsp*^F/F^:*Ctnnb1*^LoF^ and *Myh6-Mcm*^Tam^:*Dsp*^F/F^:*Ctnnb1*^GoF^. The phenotype in the *Myh6-Mcm*^Tam^, *Myh6-Mcm*^Tam^:*Dsp*^F/F^*, Myh6-Mcm*^Tam^:*Ctnnb1*^LoF^ and *Myh6-Mcm*^Tam^:*Ctnnb1*^GoF^ have been published. (11, 25, 33) Therefore, for clarity and to avoid redundancy, only the four main groups of WT, *Myh6-Mcm*^Tam^:*Dsp*^F/F^, *Myh6-Mcm*^Tam^:*Dsp*^F/F^:*Ctnnb1*^LoF^ and *Myh6-Mcm*^Tam^:*Dsp*^F/F^:*Ctnnb1*^GoF^ are included in the present study.

### Dysregulation of the cWNT pathway in the *Myh6-Mcm*^Tam^:*Dsp*^F/F^ cardiac myocytes

A total of 12,762 genes were differentially expressed in the *Myh6-Mcm*^Tam^:*Dsp*^F/F^ as compared to the WT cardiac myocytes, and were comprised of over 10,000 protein-coding and >1,000 genes coding for the long coding RNAs. (11) Three hundred forty-six genes whose transcript levels were affected by the transient expression and activation of Cre recombinase and tamoxifen injection were removed from the dataset. (33) Analysis of the remaining DEGs showed enrichment for the cWNT (TCF7L2/β-catenin) target genes in the *Myh6-Mcm*^Tam^:*Dsp*^F/F^ myocytes, comprised of 214 upregulated and 68 downregulated genes (Figure 1, A and B). The dysregulated cWNT target genes predicted the activation of EMT, and inflammatory pathways, including TNFα, TP53, and apoptosis, among others (Figure 1C).

The expression level of β-catenin (CTNNB1), a key protein in the cWNT pathway was increased, whereas the transcript level of the *Ctnnb1* gene, was unchanged in cardiac myocytes isolated from the *Myh6-Mcm*^Tam^:*Dsp*^F/F^ mouse hearts, indicating a post-transcriptional regulation (Figure 1, D-M). In addition, TCF7L2, a co-transcriptional regulator of the cWNT pathway, AXIN2, the bona fide target of the pathway, and MYC were alternatively spliced. (Figure 1, panels D-M) The aberrant isoform of each protein was upregulated, whereas the main isoforms were unchanged. (Figure 1, panels D-M) Immunofluorescence staining of thin myocardial sections corroborated increased levels of β-catenin at the IDs and increased nuclear localization of TCF7L2 in cardiac myocytes (Figure 1, panels N-R). An incidental finding was a reduced number of the myocardial cells that expressed PCM1, mainly cardiac myocytes, in the *Myh6-Mcm*^Tam^:*Dsp*^F/F^ mouse hearts (Figure 1, panels N and Q).

A notable finding in the RNA-Seq dataset was altered transcript levels of 73 inhibitors/suppressors of the cWNT/β-catenin pathway (46 upregulated and 27 suppressed) in the cardiac myocytes isolated from the *Myh6-Mcm*^Tam^:*Dsp*^F/F^ mice (Figure 2A). Genes encoding secreted cWNT inhibitors *Sfrp2*, *Dkk2*, *Tsku*, *Apoe*, and *Sost* were upregulated whereas *Wnt5b*, *Wif1*, and *Fgf9*, among others, were downregulated (Figure 2A). Increased transcript levels of selected inhibitors of the cWNT were corroborated in independent RNA samples by RT-PCR (Figure 2B). In addition, levels of DKK2, APOE, and LGALS3 proteins were increased in immunoblots in the myocardial protein extracts from the *Myh6-Mcm*^Tam^:*Dsp*^F/F^ mice (Figure 2, C and D). Similarly, immunocytochemical staining of thin myocardial sections showed increased levels of SFRP4, a classic secreted cWNT inhibitor (Figure 2E).

**Figure 2.**
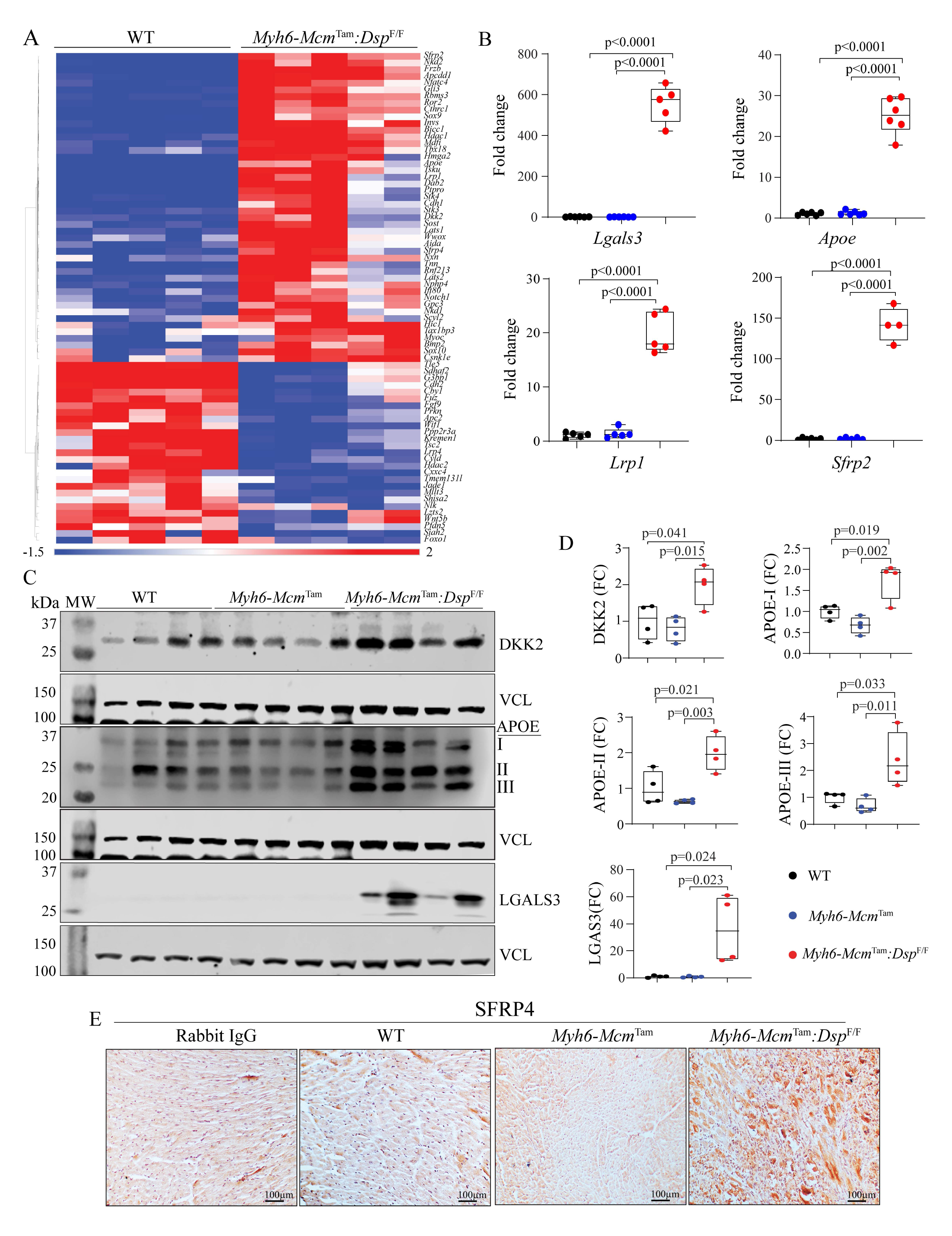
Upregulation of the cWNT pathway inhibitors in the *Myh6-Mcm*^Tam^*:Dsp*^F/F^ cardiac myocytes and myocardium. **A.** Heat map of the transcript levels of genes known to encode inhibitors of the cWNT pathway that are differentially expressed in the *Myh6-Mcm*^Tam^*:Dsp*^F/F^ cardiac myocytes. **B.** Transcript levels of selected inhibitors of the cWNT determined by RT-PCR in independent samples. **C** and **D**. Immunoblots showing protein levels of selected cWNT inhibitors and the corresponding quantitative data. **E.** Immunohistochemistry panels showing expression of SFRP4, a classic cWNT inhibitor, in the experimental groups. * denotes p<0.05 compared to WT, # p<0.05 compared to *Myh6-Mcm*^Tam^*:Dsp*^F/F^ and ¶ p<0.05 compared to *Myh6-Mcm*^Tam^*:Dsp*^F/F^*:Ctnnb1*^LoF^.

### Genetic activation or inactivation of β-catenin in the cardiac myocytes in the *Myh6-Mcm*^Tam^:*Dsp*^F/F^ mice

To discern the role of the dysregulated cWNT/β-catenin pathway in the pathogenesis of desmoplakin cardiomyopathy, β-catenin was genetically activated or inactivated in the myocytes in the *Myh6-Mcm*^Tam^:*Dsp*^F/F^ mice (Online Figure 1). Immunoblotting showed increased β-catenin levels in the *Myh6-Mcm*^Tam^:*Dsp*^F/F^ as compared to the WT mice, which was also shown in Figure 1 (Figure 3, A and B). The β-catenin level was reduced in the *Myh6-Mcm*^Tam^:*Dsp*^F/F^:*Ctnnb1*^LoF^ as compared to the *Myh6-Mcm*^Tam^:*Dsp*^F/F^ but was higher than the levels in the WT cardiac myocytes (Figure 3, A and B). The *Myh6-Mcm*^Tam^:*Dsp*^F/F^:*Ctnnb1*^GoF^ mouse cardiac myocytes showed reduced levels of the full-length β-catenin but increased levels of the ∼ 75 kDa truncated β-catenin (Figure 3, A and B). The former is likely due to incomplete Cre-mediated excision and the latter is the consequence of the deletion of exon 3 and removal of 80 amino acids (Figure 3, A and B). Immunofluorescence staining of thin myocardial sections corroborated the findings by showing increased areas stained for β-catenin expression in *Myh6-Mcm*^Tam^:*Dsp*^F/F^ mouse hearts as compared to the WT mice, reduced areas in the *Myh6-Mcm*^Tam^:*Dsp*^F/F^:*Ctnnb1*^LoF^ and increased β-catenin stained areas in the *Myh6-Mcm*^Tam^:*Dsp*^F/F^:*Ctnnb1*^GoF^ mouse hearts (Figure 3, C and D).

**Figure 3.**
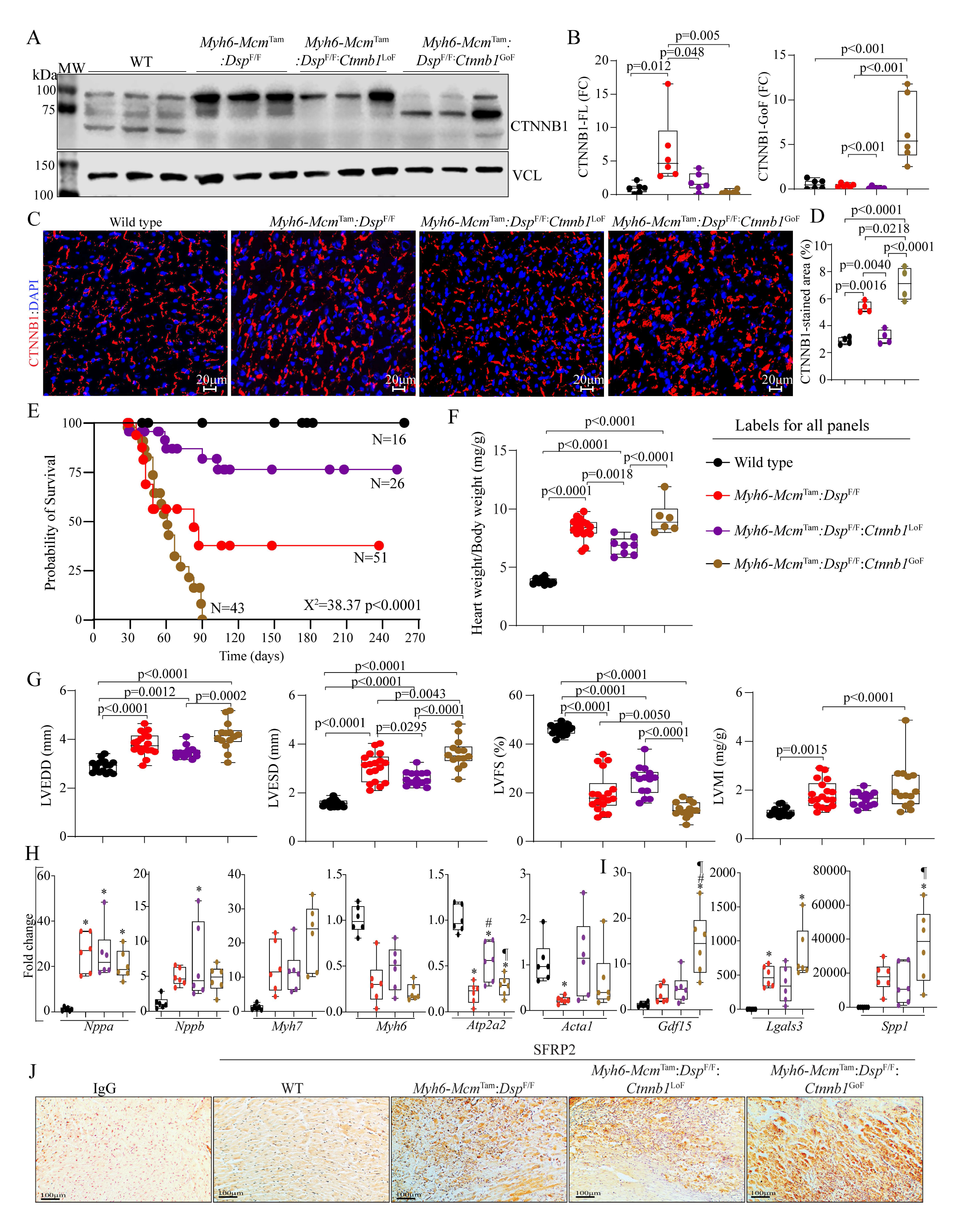
Effects of genetic inactivation and activation of β-catenin on survival and cardiac function. **A.** Immunoblot showing β-catenin expression in the wild type (WT), *Myh6-Mcm*^Tam^*:Dsp^F/F^*, *Myh6-Mcm*^Tam^*:Dsp^F/F^:Ctnnb1*^LoF^ and *Myh6-Mcm*^Tam^*:Dsp^F/F^:Ctnnb1*^GoF^ myocytes along with the vinculin (VCL), the latter as a control for loading conditions. **B.** Quantitative data showing levels of full-length and stable β-catenin (exon 3 deleted, GoF) protein. **C** and **D**. Immunofluorescence panels show expression and localization of the β-catenin in the myocardial sections along with the corresponding quantitative data. **E**. Kaplan-Meier survival plots. **F**. Heart weight/body weight ratio. **G**. Selected echocardiographic indices, namely left ventricular end-diastolic diameter (LVEDD), LV end-systolic diameter (LVESD), LV fractional shortening (LVFS), and LV mass index (LVMI). **H**. Transcript levels of selected markers of hypertrophy and failure, as analyzed by RT-PCR. **I**. Transcript levels of selected secreted biomarkers of heart failure, as determined by RT-PCR. **J.** Immunohistochemistry panels showing expression of SFRP2. Myocardial sections stained with IgG alone served as controls.

### Survival

The *Myh6-Mcm*^Tam^:*Dsp*^F/F^ mice die prematurely with a median survival time of approximately 90 days, as published (Figure 3E). (11) Genetic inactivation of β-catenin markedly prolonged survival in the *Myh6-Mcm*^Tam^:*Dsp*^F/F^ mice, as ∼ 75% of the *Myh6-Mcm*^Tam^*:Dsp*^F/F^:*Ctnnb1*^LoF^ mice survived up to 9 months (Figure 3E). In contrast, activation of β-catenin markedly reduced survival, as all *Myh6-Mcm*^Tam^*:Dsp*^F/F^:*Ctnnb1*^GoF^ mice died within 3 months of age (Figure 3E). As published, the *Myh6-Mcm*^Tam^, *Myh6-Mcm*^Tam^:*Ctnnb1*^LoF^ and *Myh6-Mcm*^Tam^:*Ctnnb1*^GoF^ mice survived normally. (25, 33)

### HW/BW

The HW/BW ratio, calculated in 4-week old mice, was increased in the *Myh6-Mcm*^Tam^:*Dsp*^F/F^, as compared to the WT mice. It remained unchanged in the *Myh6-Mcm*^Tam^*:Dsp*^F/F^:*Ctnnb1*^LoF^ and *Myh6-Mcm*^Tam^*:Dsp*^F/F^:*Ctnnb1*^GoF^ mice as compared to the *Myh6-Mcm*^Tam^:*Dsp*^F/F^ but remained increased as compared to the WT mice (Figure 3F).

### Cardiac size and function

The effects of LoF and GoF of β-catenin on cardiac size and function in the *Myh6-Mcm*^Tam^*:Dsp*^F/F^ mice were determined at 4 weeks of age by echocardiography. As reported before, deletion of *Dsp* was associated with severe cardiac dilatation and dysfunction consistent with the high mortality of the *Myh6-Mcm*^Tam^*:Dsp*^F/F^ (Figure 3G and Table 1). (11) Genetic inactivation of β-catenin in the *Myh6-Mcm*^Tam^*:Dsp*^F/F^ mice was associated with the attenuation of cardiac dilatation and dysfunction, whereas its activation has the opposite effects (Figure 3G and Table 1). The LVM index (LVMI) remained increased in all of the group with the deletion of the *Dsp* gene, compared to the WT mice (Figure 3G, Table 1).

Cardiac dysfunction is associated with the altered expression of several genes, including those whose protein products are commonly used as biomarkers to assess the severity of heart failure. Therefore, the effects of LoF and GoF of the β-catenin on selected molecular markers of hypertrophy and failure in isolated cardiac myocytes were determined by RT-PCR. By and large, the LoF and GoF of β-catenin had no significant effects on the transcript levels of the conventionally used molecular markers except *Atp2a2*, which was reduced in the *Myh6-Mcm*^Tam^*:Dsp*^F/F^ mice, it was partially rescued in the *Myh6-Mcm*^Tam^*:Dsp*^F/F^:*Ctnnb1*^LoF^, and remained suppressed in the *Myh6-Mcm*^Tam^*:Dsp*^F/F^:*Ctnnb1*^GoF^ mouse hearts (Figure 3H). Similarly, transcript levels of the selected secreted biomarkers of heart failure, namely *Gdf15*, *Lgals3*, and *Spp1* were elevated in the *Myh6-Mcm*^Tam^*:Dsp*^F/F^ mouse hearts and remained largely unaffected by the LoF and GoF of β-catenin (Figure 3, panel I). Finally, immunohistochemical staining of thin myocardial sections showed increased levels of the SFRP2 in the *Myh6-Mcm*^Tam^*:Dsp*^F/F^ mouse myocardium, which remained increased in the myocardium of DSP-deficient mice with the LoF and GoF of β-catenin (Figure 3, panel J).

### Myocyte size and number

Given the presence of cardiac dilatation and dysfunction, cardiac myocyte size, and number were assessed upon co-staining of thin myocardial sections with WGA and anti-PCM1 antibody. (25, 32, 35, 39) The myocyte CSA was increased in the *Myh6-Mcm*^Tam^*:Dsp*^F/F^ mouse hearts (Figure 4, A and B). It was normalized in the *Myh6-Mcm*^Tam^*:Dsp*^F/F^:*Ctnnb1*^LoF^, and further increased in the *Myh6-Mcm*^Tam^*:Dsp*^F/F^:*Ctnnb1*^GoF^ mouse hearts (Figure 4, A and B). The number of cells expressing PCM1, a marker mainly for cardiac myocytes in the myocardium, was reduced in the *Myh6-Mcm*^Tam^*:Dsp*^F/F^ mouse heart, was normalized in *Myh6-Mcm*^Tam^*:Dsp*^F/F^:*Ctnnb1*^LoF^ and remained reduced in the *Myh6-Mcm*^Tam^*:Dsp*^F/F^:*Ctnnb1*^GoF^ mouse hearts (Figure 4, A and C).

**Figure 4.**
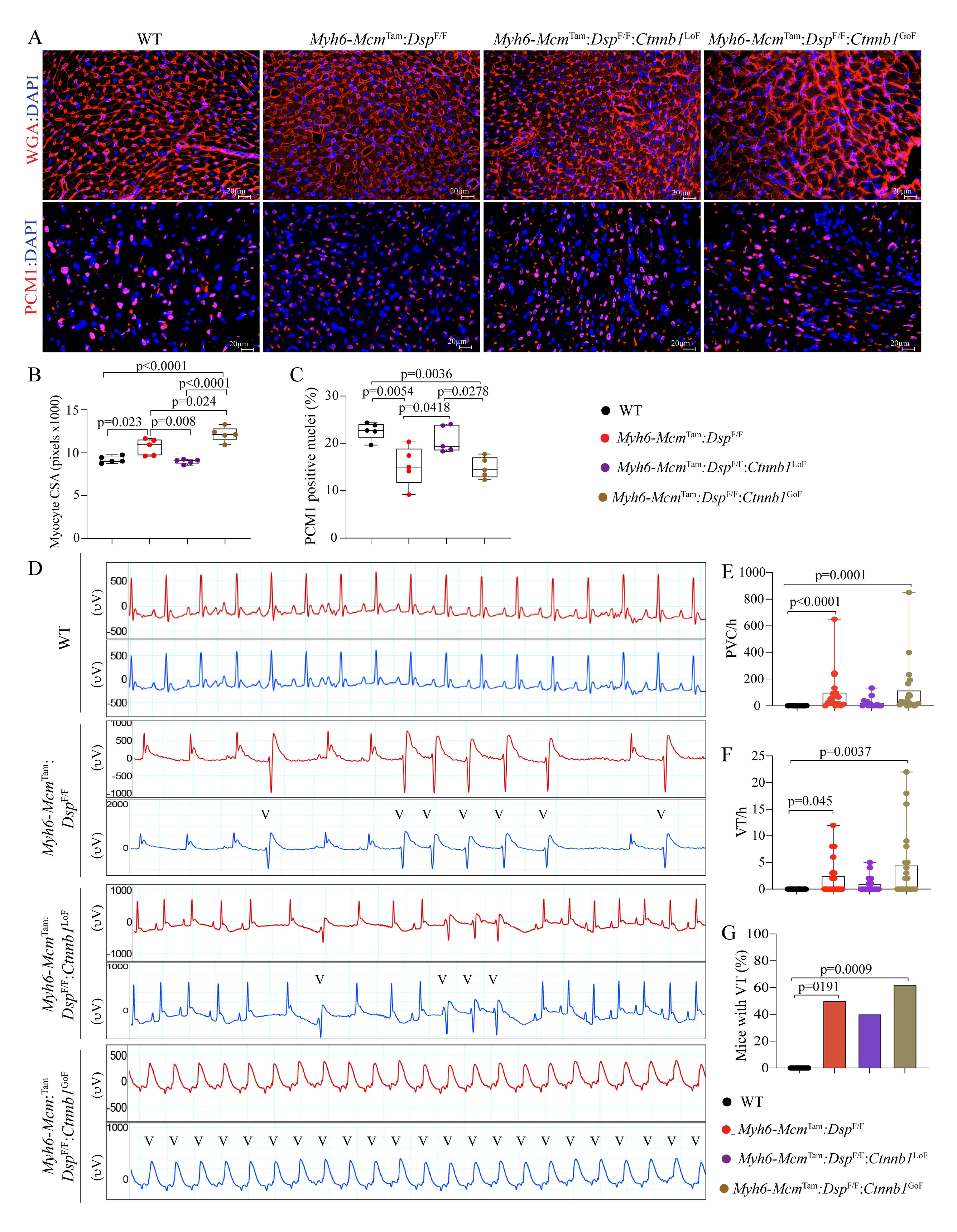
Effects of genetic inactivation and activation of β-catenin on cardiac myocyte size and arrhythmias.

**A.** Wheat Germ Agglutinin (WGA) and PCM1 co-stained thin myocardial sections were used to calculate myocyte cross-sectional area, which is depicted in panel **B.** Quantitative data depicting myocyte cross-sectional area (CSA) calculated from the WGA-stained sections and corrected for the number of myocytes. **C.** The number of myocardial cells expressing PCM1, a marker mainly of cardiac myocytes in the heart. Panel **D** illustrates selected 2-lead electrocardiograms showing the presence of ventricular ectopic beats in the experimental groups and their absence in the wild-type mice. Panels **E-G**. Quantitative data depict an increased number of premature ventricular beats (PVCs), ventricular tachycardia (VT), and the number of mice exhibiting VT in the experimental but not the control groups.

### Cardiac arrhythmias

Deletion of the *Dsp* gene in cardiac myocytes was associated with a significant increase in the prevalence of ventricular arrhythmias (Figure 4, D and E). Premature ventricular contractions (PVCs) were the most common ventricular arrhythmias and the number of PVCs per hour was increased by ∼ 15-fold in the *Myh6-Mcm*^Tam^*:Dsp*^F/F^ mice, albeit there was a wide range of variability (Figure 4, D and E). Likewise, the prevalence of ventricular tachycardia, defined as ≥ 3 PVCs was increased markedly in the *Myh6-Mcm*^Tam^*:Dsp*^F/F^ and approximately 40% of these mice exhibited VT episodes during rhythm monitoring (Figure 4, D, F, and G). The prevalence of ventricular arrhythmias trended lower in the *Myh6-Mcm:Dsp*^F/F^:*Ctnnb1*^LoF^, whereas it was the highest in the *Myh6-Mcm:Dsp*^F/F^*:Ctnnb1*^GoF^ mice, albeit the differences were confounded by a large variability in the prevalence of arrhythmias (Figure 4, D-G).

### Myocardial fibrosis

The *Myh6-Mcm*^Tam^*:Dsp*^F/F^ mice exhibit severe myocardial fibrosis, in agreement with the phenotypic feature of DC, as published. (11) Myocardial fibrosis was markedly attenuated in the *Myh6-Mcm*^Tam^*:Dsp*^F/F^:*Ctnnb1*^LoF^ and exacerbated in the *Myh6-Mcm*^Tam^*:Dsp*^F/F^:*Ctnnb1*^GoF^ mice, as determined by multiple complementary methods (Figure 5). For example, CVF comprised ∼ 25% of the myocardium in the *Myh6-Mcm*^Tam^*:Dsp*^F/F^ mice, it was reduced to ∼ 15% in the *Myh6-Mcm*^Tam^*:Dsp*^F/F^:*Ctnnb1*^LoF^ and increased to ∼ 35% in the *Myh6-Mcm*^Tam^*:Dsp*^F/F^:*Ctnnb1*^GoF^ mice (Figure 5, panels A-C). Immunostaining of thin myocardial sections for the expression of COL1A1, a major myocardial collagen, showed a similar pattern (Figure 5D). Moreover, immunoblotting of myocardial protein extracts probed with an anti-TGF β1 antibody showed increased levels of latent and mature TGFβ1 protein in the *Myh6-Mcm*^Tam^*:Dsp*^F/F^ and *Myh6-Mcm*^Tam^*:Dsp*^F/F^:*Ctnnb1*^LoF^ mice as compared to the WT mice and their further augmentations in the *Myh6-Mcm*^Tam^*:Dsp*^F/F^:*Ctnnb1*^GoF^ mice (Figure 5, E and F).

**Figure 5.**
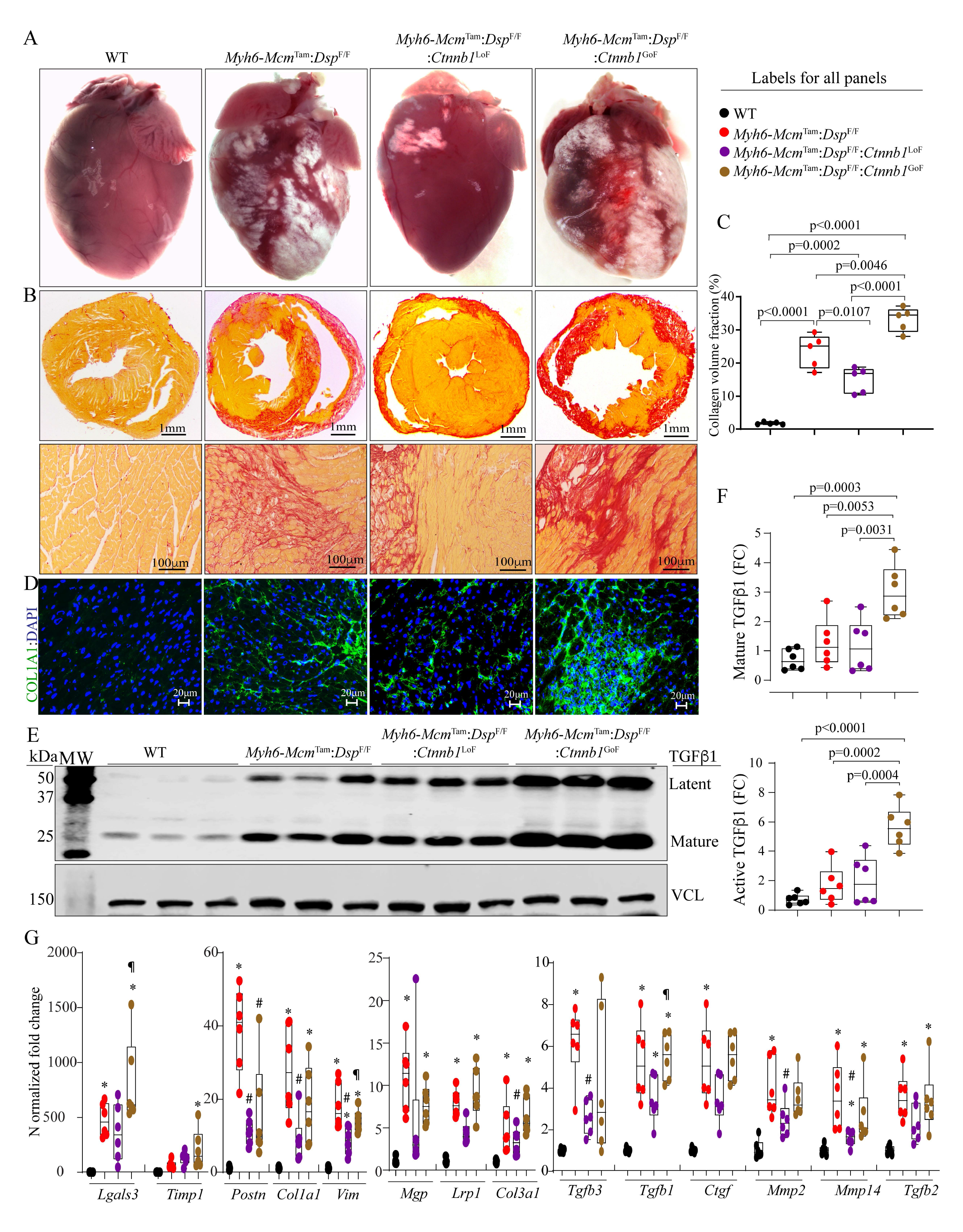
Effects of genetic inactivation and activation of β-catenin on myocardial fibrosis. Panel **A** exhibits gross heart morphology, showing extensive epicardial fibrosis (white areas) in the experimental groups. Panel **B** shows picrosirius red stained thin myocardial cross-sections (upper panels) and high magnification thin myocardial sections (lower panels), used to calculate collagen volume fraction (CVF). **C**. Graph depicts CVF in the study groups. **D.** Immunofluorescence panels showing myocardial sections stained with an antibody against COL1A1. **E.** and **F**. Immunoblots showing expression of TGFβ1, latent (upper panel), and active (lower panel) in the study groups along with the quantitative data for each isoform. **G**. Transcript levels of selected genes involved in fibrosis as quantified by RT-PCR. Symbols as in Figure 4.

Transcript levels of several genes involved in myocardial fibrosis were analyzed by RT-PCR in cardiac RNA extracts, which showed marked increases in the levels of *Tgfb1*, *Tgfb2*, *Tgfb3*, *Postn*, *Pdgfra*, *Col1a1*, *Col1a3*, *Ctgf*, *Mmp2*, *Mmp14*, *Lrp1*, *Mgp*, *Vim*, *Timp1*, and *Lgals3* in *Myh6-Mcm*^Tam^*:Dsp*^F/F^ mouse hearts and the pattern of attenuation or augmentation in the *Myh6-Mcm*^Tam^*:Dsp*^F/F^:*Ctnnb1*^LoF^ and *Myh6-Mcm*^Tam^*:Dsp*^F/F^:*Ctnnb1*^GoF^ mice, respectively (Figure 5G).

### PANoptosis

PANoptosis, comprised of apoptosis, necroptosis, and pyroptosis, is a prominent phenotypic feature of the mouse model of DC in agreement with the prominence of cell death in human DC. (7, 11) Thus, selected markers of PANoptosis were analyzed to assess the effects of LoF and GoF of the β-catenin in the *Myh6-Mcm*^Tam^*:Dsp*^F/F^ mice. TUNEL assay showed an increased number of cells stained positive in the *Myh6-Mcm*^Tam^*:Dsp*^F/F^ mice, an attenuated number in the *Myh6-Mcm*^Tam^*:Dsp*^F/F^:*Ctnnb1*^LoF^ and an increased number in the *Myh6-Mcm*^Tam^*:Dsp*^F/F^:*Ctnnb1*^GoF^ mouse hearts (Figure 6, A and B). Immunoblot analysis of myocardial protein extracts showed increased levels of several proteins involved in apoptosis, necroptosis, and pyroptosis, namely CASP3, RIPK1 and 3, MLKL, GSDMD, and ASC in the *Myh6-Mcm*^Tam^*:Dsp*^F/F^ mouse hearts, consistent with the previous data. (11) The LoF of β-catenin attenuated but not normalized upregulated expression levels of the above proteins, whereas the GoF of β-catenin further augmented the upregulated proteins involved in selected cell death programs (Figure 6, C and D). The pattern of attenuation upon the LoF of β-catenin and augmented expression upon the gain of function of β-catenin was consistent among the selected proteins.

**Figure 6.**
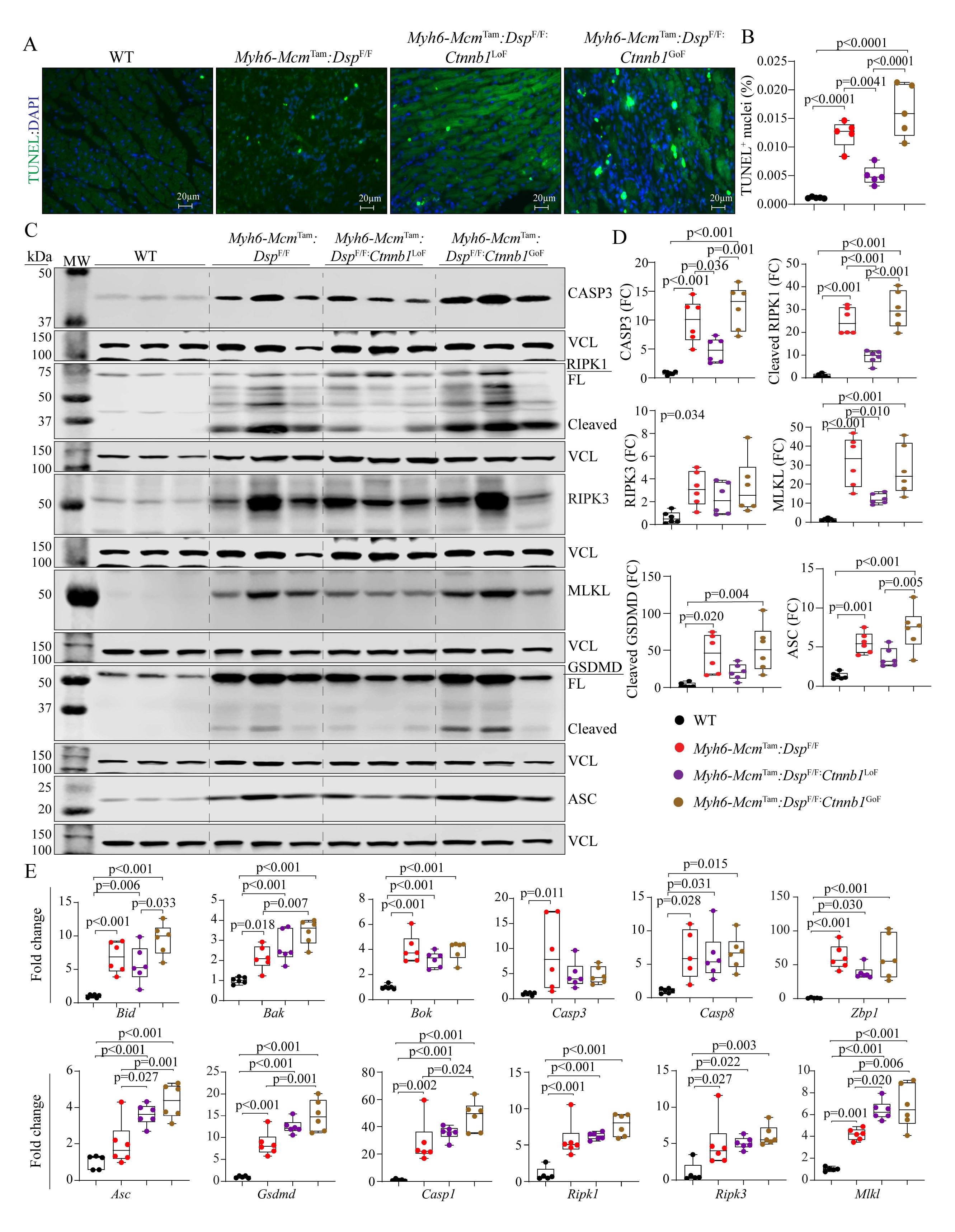
Effects of genetic inactivation and activation of β-catenin on PANoptosis. Panel **A** shows thin myocardial sections stained with the TUNEL assay and DAPI, the latter marks the DNA. B. Quantitative data showing the percentage of myocardial cells stained positive for the TUNEL assay. **C** and **D**. Immunoblot panels showing expression of selected proteins involved in PANoptosis and the corresponding quantitative data. CASP3 is used as a marker of apoptosis, RIPK1, RIPK3, and MLKL are used as markers of necroptosis, and GSDMD and ASC as markers of pyroptosis. Panel **E** shows transcript levels of a dozen genes involved in cell death programs, as determined by RT-PCR.

Analysis of the transcript levels of a dozen genes involved in PANoptosis in isolated cardiac myocytes showed markedly increased levels in the *Myh6-Mcm*^Tam^*:Dsp*^F/F^ mice. However, the LoF and GoF of β-catenin, did not have consistent effects on the transcript levels, partly reflective of regulation of the expression levels of the selected markers at the post-transcriptional level, albeit transcript levels of several genes, such as *Bak*, *Casp1*, *Mlkl,* and *Asc*, were further upregulated in the GoF group (Figure 6E).

### Desmosome protein expression and localization

The desmosomes undergo extensive remodeling in ACM, including in the *Myh6-Mcm*^Tam^*:Dsp*^F/F^ mice, evidenced by decreased levels of several key protein constituents of desmosomes and altered desmosome structures. (11, 15, 44) To determine whether the LoF and GoF of β-catenin affected expression levels and localization of selected desmosome proteins, myocardial protein extracts were probed, and thin myocardial sections were stained with antibodies against the selected proteins. As reported previously, JUP, DSG2, DSC2, and PKP2 proteins were reduced in the *Myh6-Mcm*^Tam^*:Dsp*^F/F^ mouse hearts, and remained largely suppressed in the *Myh6-Mcm*^Tam^*:Dsp*^F/F^:*Ctnnb1*^LoF^ and *Myh6-Mcm*^Tam^*:Dsp*^F/F^:*Ctnnb1*^GoF^ mouse hearts, except for a modest increase in the JUP and PKP2 expressions in the LoF and further suppression in the GoF of β-catenin groups (Online Figure 2, A and B). The findings of immunofluorescence staining of myocardial sections were largely similar to the immunoblotting data (Online Figure 2C).

### Transcriptional and biological pathways

To gain insights into the mechanism(s) by which the LoF of β-catenin attenuates and the GoF exacerbates the phenotype in the *Myh6-Mcm*^Tam^*:Dsp*^F/F^ mice, transcripts of isolated cardiac myocytes were analyzed by RNA-Seq. The data were analyzed after the removal of 346 genes whose expressions were affected by tamoxifen injection and expression and activation of Cre recombinase, as defined previously. (33) Some of the quality control metrics of the RNA-Seq data are shown in Online Table 2 and those in the *Myh6-Mcm*^Tam^ alone, *Myh6-Mcm*^Tam^:*Dsp*^F/F^*, Myh6-Mcm*^Tam^:*Ctnnb1*^LoF^, and *Myh6-Mcm*^Tam^:*Ctnnb1*^GoF^ mice have been published. (25, 33)

To determine whether phenotypic effects of LoF and GoF of β-catenin were reflective of changes in the transcriptional activity of the cWNT pathway, transcript levels of the recently defined cWNT target genes in the mouse cardiac myocytes, were compared among the groups. (25) Only a small fraction of the 1,075 cWNT target genes in the mouse cardiac myocytes were changed upon LoF (15 genes) or GoF (144 genes) of β-catenin (Figure 7A). Comparing the cardiac myocyte transcripts between the *Myh6-Mcm*^Tam^:*Dsp*^F/F^:*Ctnnb1*^LoF^ and *Myh6-Mcm*^Tam^:*Dsp*^F/F^:*Ctnnb1*^GoF^ showed differential expression of only 46 cWNT target genes, comprised of 13 upregulated and 33 downregulated genes in the β-catenin LoF genotype (Figure 7A).

**Figure 7.**
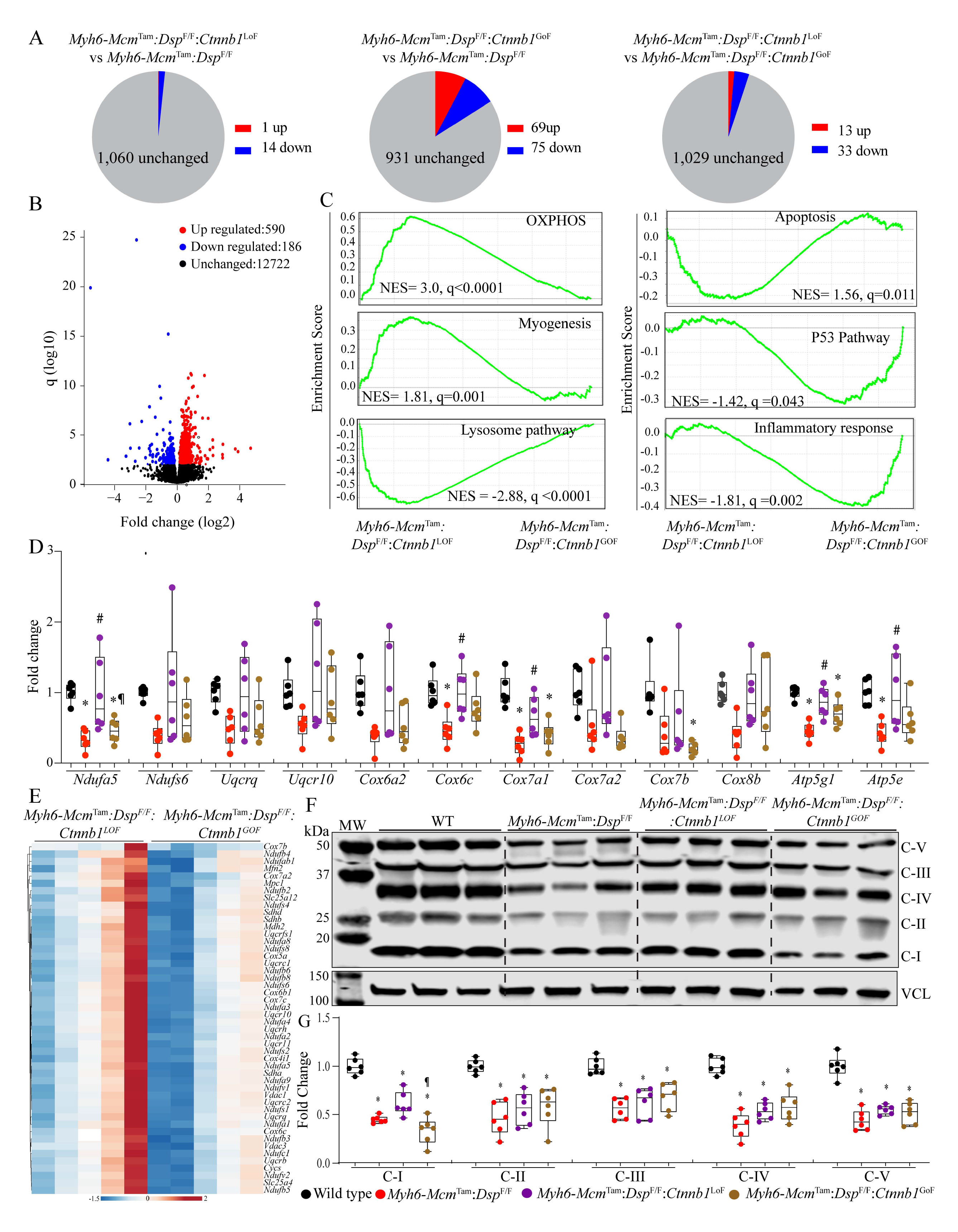
Effects of genetic inactivation and activation of β-catenin on biological pathways. Panel **A** illustrates pie charts showing the number of the cWNT target genes in cardiac myocyte RNA-Seq data, which were affected by the genetic inactivation and activation of β-catenin in the *Myh6-Mcm*^Tam^:*Dsp*^F/F^ mice. The cWNT target genes used in this analysis were obtained from the previously published list of genes that were induced in cardiac myocytes upon activation of the β-catenin. (25) **B**. Volcano plot showing differentially expressed genes between the *Myh6-Mcm*^Tam^*:Dsp*^F/F^*:Ctnnb1*^LoF^ and *Myh6-Mcm*^Tam^*:Dsp*^F/F^*:Ctnnb1*^GoF^ myocytes. **C**. GSEA plots showing biological pathways (Hallmark signature) that were predicted to be differential between the *Myh6-Mcm*^Tam^*:Dsp*^F/F^*:Ctnnb1*^LoF^ and *Myh6-Mcm*^Tam^*:Dsp*^F/F^*:Ctnnb1*^GoF^ myocytes. **D**. Data illustrates transcript levels of selected genes involved in oxidative phosphorylation (OXPHOS) in independent RNA extracts, as determined by RT-PCR. **E**. Heat map of genes involved in OXPHOS, which were shown to be enriched in the Hallmark OXPHOS gene signature and were differentially expressed between the two groups. **F.** shows an OXPHOS blot of one selected protein in each of the 5 mitochondrial OXPHOS complexes detected using an antibody cocktail. Complex I is represented by NDUFB8, complex II by SDHB, complex III by UQCRC2, complex IV by MTCO1, and complex V by ATP5A. **G.** Quantitative data of each OXPHOS protein in panel F, normalized to vinculin (VCL). Symbols as in Figure 4.

To identify the DEGs, regardless of the cWNT targets, cardiac myocyte transcripts were compared between the *Myh6-Mcm*^Tam^:*Dsp*^F/F^ and *Myh6-Mcm*^Tam^:*Dsp*^F/F^:*Ctnnb1*^LoF^, which showed differential expression of 205 genes (Online Figure 3, A and B). The DEGs predicted increased activities of transcriptional regulators TP53 and NCOA1 and reduced activities of MYC, TP63, RB1, MYCN, and TP73 in the *Myh6-Mcm*^Tam^:*Dsp*^F/F^:*Ctnnb1*^LoF^ as compared to *Myh6-Mcm*^Tam^:*Dsp*^F/F^ myocytes (Online Figure 3C). Given the relatively small number of DEGs, no further analysis was performed. A total of 2,610 genes were differentially expressed between *Myh6-Mcm*^Tam^:*Dsp*^F/F^ and *Myh6-Mcm*^Tam^:*Dsp*^F/F^:*Ctnnb1*^GoF^ myocytes (Online Figures 4, A and B). The KDM5A, TP53, BACH1, and several other transcriptional regulators were predicted to be activated in the *Myh6-Mcm*^Tam^:*Dsp*^F/F^:*Ctnnb1*^GoF^ myocytes, whereas RB1, NFE2L2, and several others were predicted to be suppressed (Online Figure 4, C and D). The GSEA predicted activation of INFγ response, EMT, and apoptosis and suppression of OXPHOS, adipogenesis, and myogenesis in the *Myh6-Mcm*^Tam^:*Dsp*^F/F^:*Ctnnb1*^GoF^ myocytes (Online Figure 4E).

To gain further insights into the pathways regulated by the β-catenin, which might contribute to the partial phenotypic rescue in the *Myh6-Mcm*^Tam^*:Dsp*^F/F^ mice, gene expression between the *Myh6-Mcm*^Tam^*:Dsp*^F/F^:*Ctnnb1*^LoF^ and *Myh6-Mcm*^Tam^*:Dsp*^F/F^:*Ctnnb1*^GoF^ were compared. There were 880 DEGs, which were comprised of 652 upregulated and 228 downregulated genes in the *Myh6-Mcm*^Tam^*:Dsp*^F/F^:*Ctnnb1*^LoF^ compared to the *Myh6-Mcm*^Tam^*:Dsp*^F/F^:*Ctnnb1*^GoF^ myocytes (Figure 7B and Online Figure 5A). The GSEA predicted suppression of the cWNT/β-catenin signaling pathway, per the fidelity of the LoF and GoF models, and suppression of transcriptional regulators TP53 and KDM5A and activation of RB1 and PPARGC1A, among others in the *Myh6-Mcm*^Tam^:*Dsp*^F/F^*:Ctnnb1*^LoF^ myocytes (Online Figure 5, B and C). The DEGs also predicted suppression of the inflammatory pathways and activation of OXPHOS and metabolic pathways in the *Myh6-Mcm*^Tam^*:Dsp*^F/F^:*Ctnnb1*^LoF^ myocytes (Online Figure 5D). Analysis of the transcripts by GSEA in the *Myh6-Mcm*^Tam^*:Dsp*^F/F^:*Ctnnb1*^LoF^ predicted enrichment of genes involved in OXPHOS, myogenesis, and suppression of the apoptosis, TP53, inflammatory responses, and lysosomal pathways, as compared to the *Myh6-Mcm*^Tam^*:Dsp*^F/F^:*Ctnnb1*^GoF^ myocytes (Figure 7C). Transcript levels of several genes involved in inflammation and lysosomal pathways were analyzed by RT-PCR in independent samples, which showed increased levels of the transcripts in the *Myh6-Mcm*^Tam^:*Dsp*^F/F^, attenuation of several but not all transcript levels in the *Myh6-Mcm*^Tam^:*Dsp*^F/F^*:Ctnnb1*^LoF^ and further upregulation in the *Myh6-Mcm*^Tam^*:Dsp*^F/F^*:Ctnnb1*^GoF^ (Online Figure 6).

Given that the GSEA and pathway analysis predicted a partial rescue of OXPHOS as one of the several putative mechanisms for the phenotypic attenuation upon LoF of β-catenin in the *Myh6-Mcm*^Tam^:*Dsp*^F/F^ mice, transcript levels of a dozen genes involved in OXPHOS in cardiac myocytes were determined in independent samples, which showed suppressed levels in the *Myh6-Mcm*^Tam^:*Dsp^F/F^* mouse myocytes as compared to the levels in the WT myocytes (Figure 7D). Likewise, a heat map of transcript levels of selected genes involved in OXPHOS in the *Myh6-Mcm*^Tam^*:Dsp*^F/F^:*Ctnnb1*^LoF^ and *Myh6-Mcm*^Tam^*:Dsp*^F/F^:*Ctnnb1*^GoF^ myocytes largely corroborated the partial rescues of genes involved in OXPHOS in the *Myh6-Mcm*^Tam^:*Dsp*^F/F^*:Ctnnb1*^LoF^ myocytes (Figure 7E). Overall, the LoF of β-catenin partially restored suppressed transcript levels of the OXPHOS genes, however, the transcript levels remained suppressed in the *Myh6-Mcm*^Tam^:*Dsp*^F/F^:*Ctnnb1*^GoF^ mice (Figure 7F). Finally, immunoblotting of selected proteins in 5 OXPHOS complexes largely corroborated the findings of RNA-Seq and RT-PCR, showing reduced levels of the selected proteins in the *Myh6-Mcm*^Tam^:*Dsp^F/F^* mouse hearts and partial rescue of OXPHOS complex I protein in the *Myh6-Mcm*^Tam^*:Dsp*^F/F^:*Ctnnb1*^LoF^, whereas levels of the selected protein in other OXPHOS complexes remained suppressed (Figure 7G and H).

## DISCUSSION

The findings are notable for the unexpected beneficial effects of reduced expression of β-catenin and the deleterious effects of increased expression of a stable β-catenin, lacking its degron, in a mouse model of ACM generated by tamoxifen-inducible deletion of the *Dsp* gene in the post-natal cardiac myocytes (DC model). The contrasting effects of LoF and GoF of β-catenin were remarkable for several phenotypes, including survival, cardiac function, arrhythmias, myocardial fibrosis, and the cell death programs, referred to as PANaptosis. Likewise, the effects on each phenotype, except for survival, were analyzed by complementary methods and the findings were concordant. The beneficial effects of the suppression of β-catenin, however, were partial, as would be expected, given the multifarity of the pathways involved in the pathogenesis of DC, as well as incomplete tamoxifen-induced Cre-mediated recombination. Likewise, the effects were judged to be independent of the role of the β-catenin in transcriptional activity of the cWNT pathway, as too few cWNT target genes in cardiac myocytes were changed upon LoF of β-catenin, which contrasts with marked phenotypic benefits. The data implicate partial restoration of the transcript and protein levels of suppressed genes involved in OXPHOS, as a putative mechanism for the salubrious effects of LoF of β-catenin.

The cWNT pathway, which is one of the most conserved pathways across the animal kingdom, is comprised of a large number of protein constituents that converge on the β-catenin/TCF7L2 transcriptional machinery to induce gene expression. (45) Whereas the majority of the previous studies have reported suppression of the cWNT pathway in ACM, data in the present study indicate dysregulation of the cWNT signaling pathway, as both the target genes of the cWNT were upregulated in the *Myh6-Mcm*^Tam^*:Dsp*^F/F^ mouse myocytes as well as concomitantly several well-established inhibitors of the cWNT pathway, including secreted cWNT inhibitors. (46) In addition, TCF7L2, the co-transcriptional effector of the cWNT pathway, and AXIN2, which is considered the bona fide target of the cWNT pathway, were alternatively spliced and the variant isoforms, with poorly defined functions, were upregulated. Thus, the data suggest a marked dysregulation of the cWNT pathway in cardiac myocytes in a mouse model of DC.

β-catenin is an indispensable and non-redundant member of the cWNT pathway. (45, 47) Consequently, it has emerged as the most effective target for genetic manipulation of the cWNT pathway. (45, 47) Pharmacological inhibition of GSK3β, which phosphorylates β-catenin at its degron sites, have been reported to improve the ACM phenotype in mouse and zebrafish models. (19-21) Likewise, pharmacological inhibition of the cWNT pathway by WNT974, which inactivates porcupine, responsible for palmitoylation and subsequent secretion of WNT ligand, exerts salutary effects. (22) The present study, by design, is distinct from the previous studies, as it utilizes complementary genetic LoF and GoF approaches to target the β-catenin. The study provides evidence at multiple levels that LoF of β-catenin is beneficial in ACM and its GoF is deleterious in a mouse model of DC. The beneficial effects seem to be independent of the transcriptional activities of the cWNT pathway.

β-catenin is not only a co-transcriptional regulator of the cWNT signaling pathway but also has several non-canonical functions, including regulation of the Hippo, SOX, FOXO, TBX5, GATA4, KLF15, NOTCH1, and HIF1α pathways. (15, 24, 48, 49) It is a major constituent of the IDs, where its C-terminal domain interacts with the cadherins and plays a crucial role in cell-cell adhesion and mechanotransduction. (17, 18) The increase in the β-catenin levels in the *Myh6-Mcm*^Tam^*:Dsp*^F/F^ mice likely reflects its junctional localization secondary to molecular remodeling of the IDs. (11) β-catenin and junction plakoglobin (JUP) are homologous proteins with a high level of amino acid identity and are partially interchangeable at the IDs and the nucleus. (23, 50, 51) Moreover, β-catenin functions are regulated by extensive post-translational modifications, including phosphorylation in over 2 dozen serine, tyrosine, threonine amino acids throughout the length of the protein by over a dozen kinases, acetylation, and ubiquitylation, among others. (26, 52) The regulatory role of the post-translational modification of β-catenin is well recognized for its flux through the AXIN-APC destruction scaffold, wherein coordinated phosphorylation of β-catenin degron by CKI and GSK3β facilitate its ubiquitylation and degradation, whenever the WNT signaling is inactive. (52) Pertinent to the present study, β-catenin is implicated in the regulation of OXPHOS, and inducible deletion of β-catenin has been shown to upregulate expression of genes encoding protein constituents of the OXPHOS, particularly the complex I proteins. (25, 53, 54) The findings of the present study implicate partial recovery of suppressed OXPHOS, as a putative mechanism for the partial phenotypic rescue in the *Myh6-Mcm*^Tam^:*Dsp*^F/F^:*Ctnnb1*^LoF^ mice. Given the scant number of altered cWNT/β-catenin target gene expressions in the *Myh6-Mcm*^Tam^:*Dsp*^F/F^:*Ctnnb1*^LoF^ as compared to the *Myh6-Mcm*^Tam^:*Dsp*^F/F^ cardiac myocytes, the partial rescue of the OXPHOS and the other phenotypes upon inactivation of the β-catenin is likely independent of the transcriptional activity of the cWNT/β-catenin pathway.

Fidelity of the genetic LoF and GoF of β-catenin in cardiac myocytes in mice has been established. (25-28) Likewise, the phenotypic consequences of short-term genetic activation or suppression of the β-catenin in mouse cardiac myocytes have been published. (25) Overall, the LoF and GoF of β-catenin had no discernible effect on survival, cardiac function, myocardial fibrosis, and cell death at 4 weeks old mice, the time point that corresponds to the mouse age in the present study. (25) The findings were corroborated using complementary methods, which showed mainly concordant results. In addition, the contrasting phenotypic directions between the GoF and LoF of the β-catenin mice provide further credence to the findings. Finally, several quality control measures were incorporated into the study design, including the removal of the genes whose expressions were affected by the expression of Cre recombinase and the administration of tamoxifen from the RNA-Seq data before additional analysis of the RNA-Seq data. (33)

The study has several limitations. Notable among them is the restriction of the findings to a single mouse model of cardiomyopathy whereby the *Dsp* gene was almost completely deleted. This contrasts with most human cardiomyopathy, which is typically caused by heterozygous for the *DSP* mutations, albeit homozygous mutations in the *DSP* and other genes have been reported in humans with ACM. (55-57) Likewise, the *Dsp* gene was deleted conditionally at post-natal day 14, in contrast to the human genotype whereby the mutation is present since the single-cell embryo formation. This approach was taken to avoid the confounding effects of activation or suppression of the cWNT/β-catenin pathway during embryogenesis, as the cWNT/β-catenin is known to affect cardiac development. (16, 31) Furthermore, *DSP* mutations cause a partially distinct phenotype notable for marked fibrosis and left ventricular dysfunction, which was also observed in the present mouse model, but the findings might not be fully applicable to ACM caused by mutations in other genes. Moreover, the phenotypic rescue upon the LoF of β-catenin in the *Myh6-Mcm*^Tam^:*Dsp*^F/F^ mice was incomplete, in part because of the multifarity of the involved mechanisms in the pathogenesis of DC and partly of partial deletion of the *Ctnnb1* gene because of incomplete recombination event and residual expression of the full size β-catenin. Furthermore, the mechanisms by β-catenin suppresses and its LoF upregulates transcription of the genes involved in OXPHOS, presumably independent of the cWNT pathway, remain to be delineated. Finally, the β-catenin is implicated in regulating energy metabolism and mitochondrial function, which were not addressed in the present study. (58)

In conclusion, the findings suggest that the genetic suppression of the β-catenin in a mouse model of DC imparts beneficial effects on survival, cardiac function, arrhythmias, myocardial fibrosis, and cell death, whereas its activation is deleterious. The findings set the stage for pilot interventional studies to explore the effects of the suppression of the β-catenin on the phenotypic expression of ACM in larger animal models and subsequently in humans.

## Figure Legends

- Only significant p values (Bonferroni corrected) are depicted in the panels throughout all figures. Each dot in the panels represents one independent sample throughout the figures
- Whenever a membrane is probed for multiple test proteins, the same blot for the loading control is included in the figures.
- The same color symbols in all quantitative panels is used for consistency and only one set of symbols is listed in each panel to avoid redundancy.

## Contributions of the authors

Melis Olcum performed most of the molecular biology and histological experiments, echocardiography, and electrocardiographic rhythm monitoring, analyzed the data, and edited the manuscript.

Siyang Fan generated and established the mouse colony, performed molecular biology experiments, echocardiography, and cardiac rhythm monitoring, and prepared the samples for RNA-Seq.

Leila Rouhi performed part of the RT-PCR, immunoblotting, and immunofluorescence studies and edited the manuscript.

Sirisha Cheedipudi performed immunofluorescent staining

Benjamin Cathcart performed part of the molecular biology experiments.

Hyun-Hwan Jeong mapped the RNA-Seq data to the mouse reference genome and performed the initial bioinformatics analysis of the RNA-Seq data.

Zhongming Zhao supervised the bioinformatics analysis of the RNA-Seq data.

Priyatansh Gurha performed the secondary analysis of the RNA-Seq data, interpreted the findings, and edited the manuscript.

AJ Marian developed the concept, supervised the experiments, interpreted the findings, and wrote the manuscript.

## Acknowledgement

Dr. Marian is supported by the National Heart, Lung, and Blood Institute (HL151737, HL132401). Dr. Gurha is supported by R56HL165334-01. The authors thank the technical support from the Cancer Genomics Core funded by the Cancer Prevention and Research Institute of Texas (CPRIT RP180734).

