## Supplements for "Genetic Inactivation of β-Catenin Attenuates and Its Activation Aggravates Desmoplakin Cardiomyopathy"

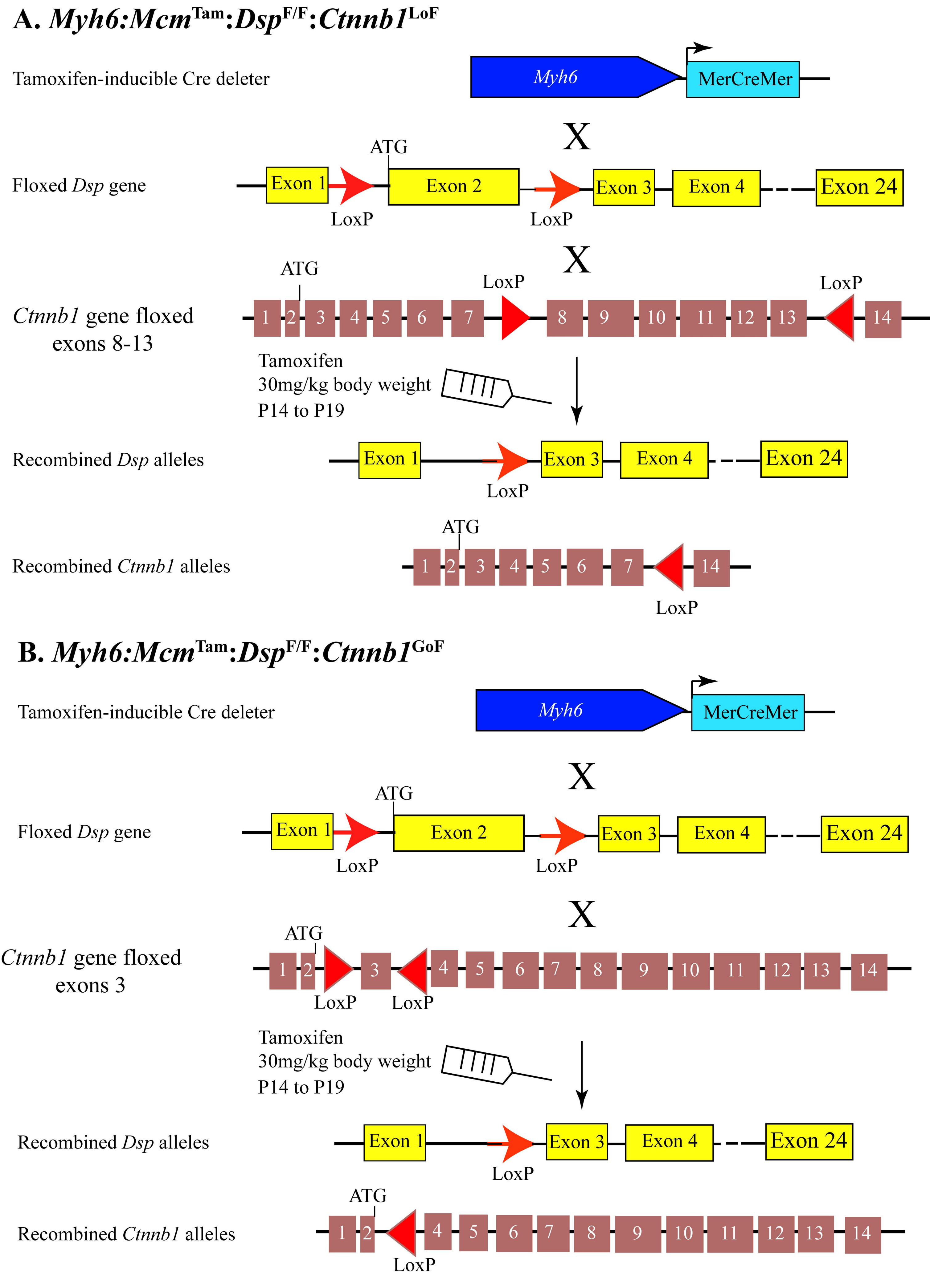


**Online Figure 1.** Schematic illustration of concomittant deletion of the *Dsp* gene along with the loss or gain of function of the ß catenin in using the *Myh6-MerCreMer, Dsp*-floxed and *Ctnnb1*-floxed alleles,

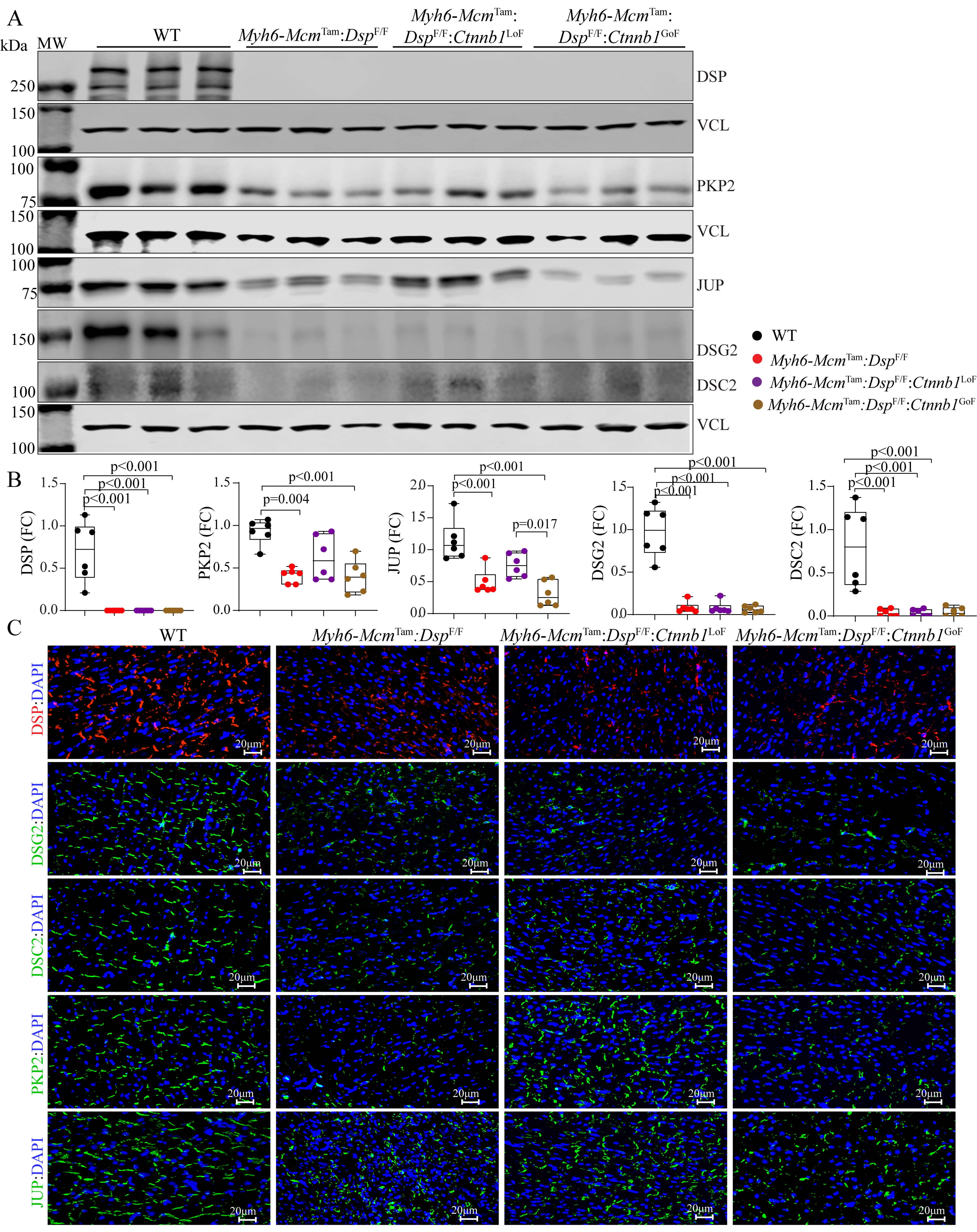


**Online Figure 2.** Effects of genetic inactivation and activation of β−catenin on expression of selected desmosome proteins. A. Immunoblots of selected desmosome proteins. B. The corresponding quantitative data of the immunoblots. C. Immunofluorescence stained myocardial sections with antibodies against selected desmosome proteins.

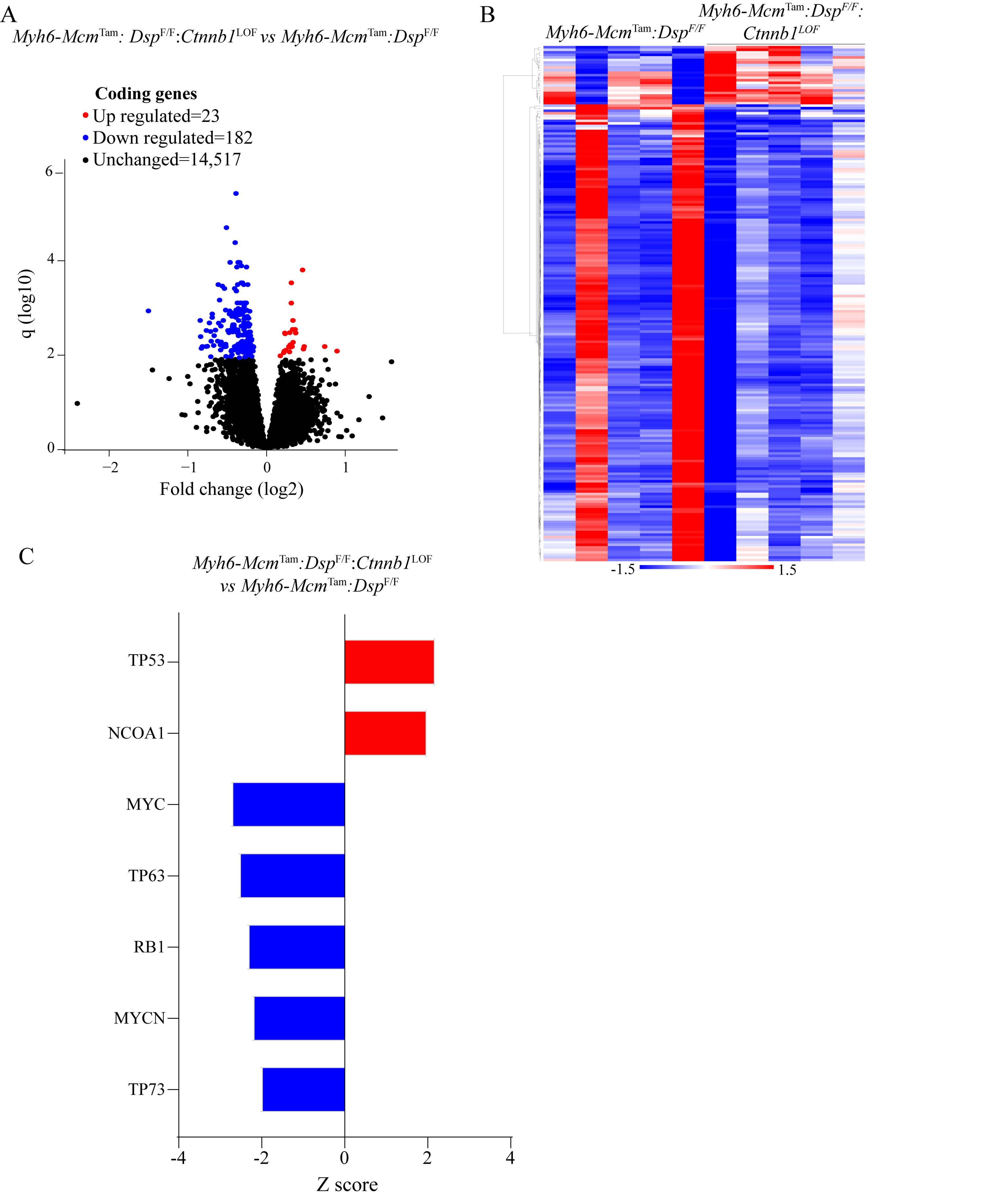


**Online Figure 3.** Changes in gene expression upon genetic inactivation of β−catenin in the *Myh6-Mcm*$^{Tam}$:*Dsp*$^{F/F}$ mouse cardiac myocytes. A. Volcano plot showing differentially expressed genes. B. Heat map of the differentially expressed genes. C. Predicted up- and down-regulated transcriptional regulators.

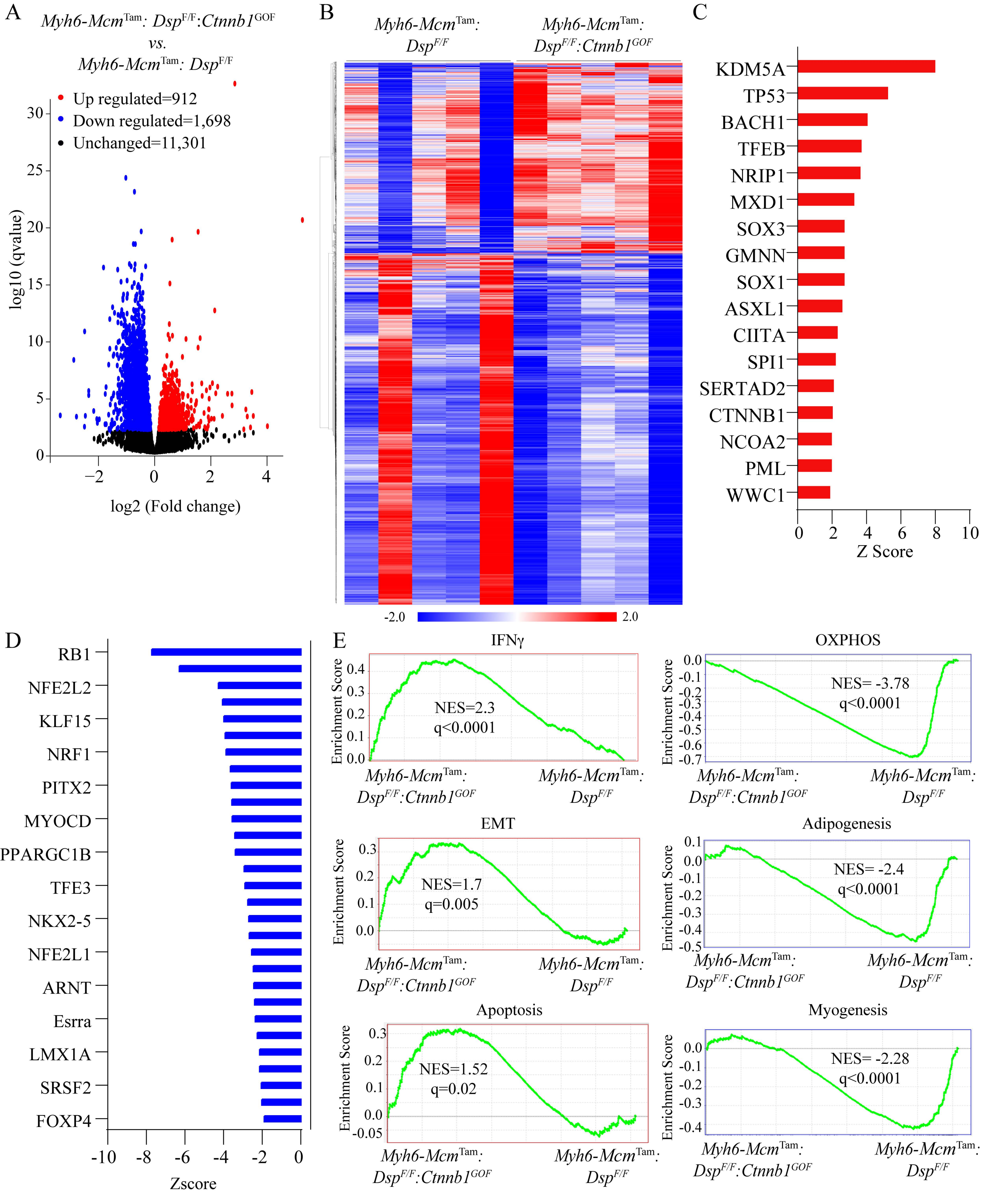


**Online Figure 4.** Effects of activation of β-catenin on gene expression in the *Myh6-Mcm*$^{Tam}$:*Dsp*$^{F/F}$ mouse myocytes. A. Volcano plot of differentially expressed genes. B. Heat map of the differentially expressed genes. C and D. Predicted dysregulated transcriptional regulators. E. GSEA plots of predicted biological pathways.

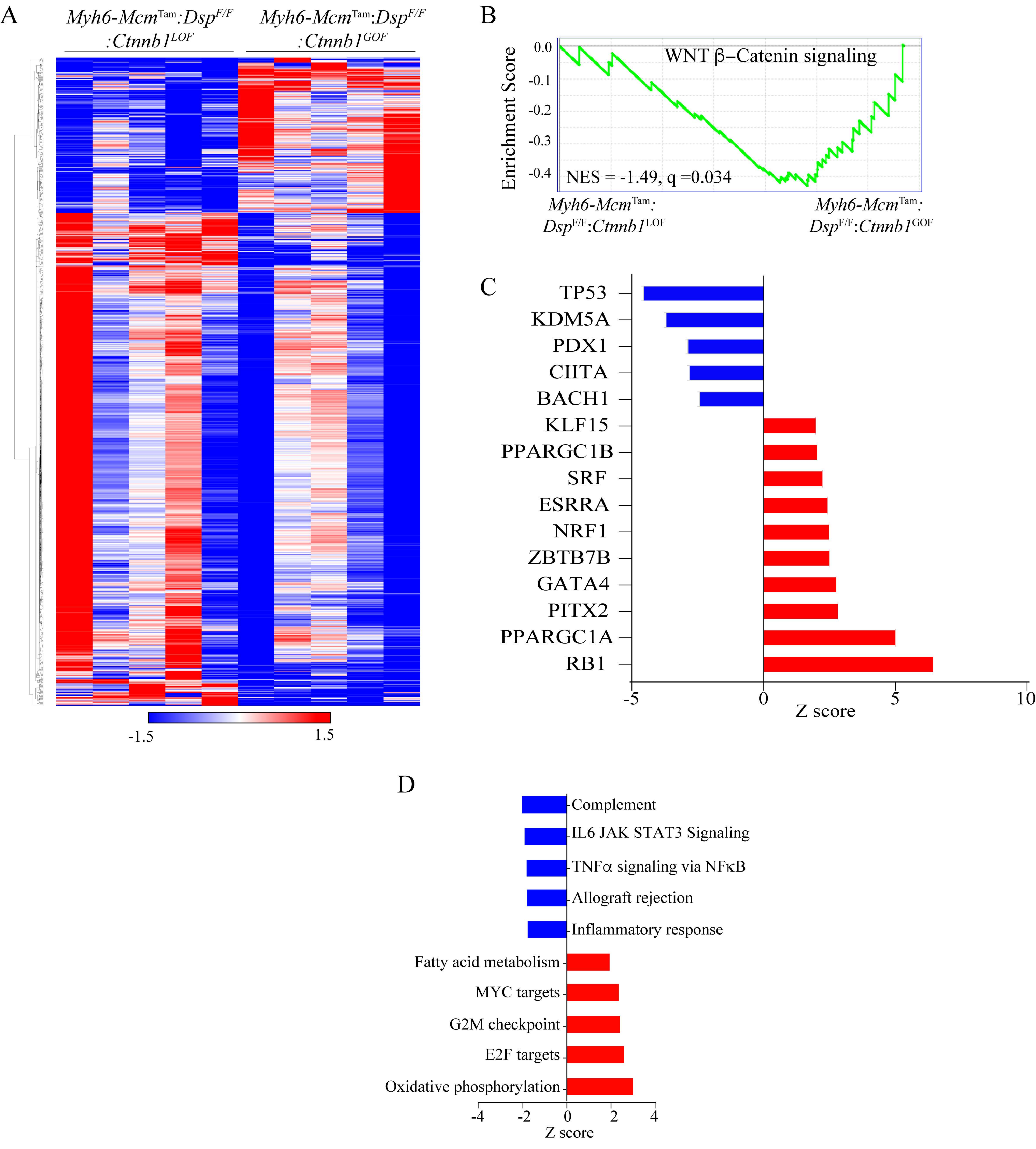


**Online Figure 5.** Comparing cardiac myocyte gene expression upon inactivation and activation of β-catenin in the *Myh6-Mcm*$^{Tam}$:*Dsp*$^{F/F}$ mice. A. Heat map of the differentially expressed genes. B. GSEA plot showing suppression of the cWNT pathway in the *Myh6-Mcm*$^{Tam}$:*Dsp*$^{F/F}$:*Ctnnb1*$^{LOF}$ myocytes. C. and D. Predicted dysregulated transcriptional regulator of gene expression and the biological pathways.

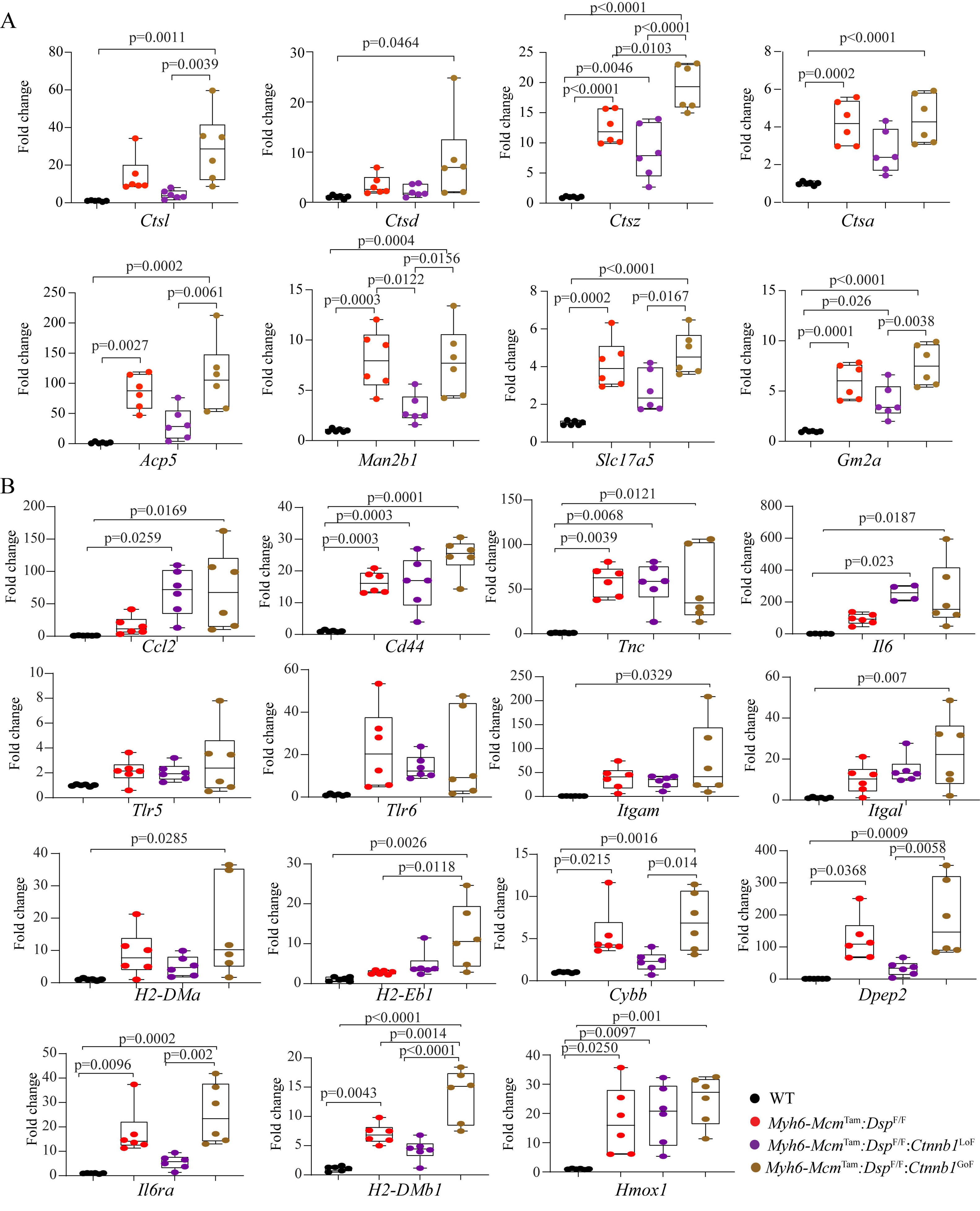


**Online Figure 6.** Transcript levels of selected genes involved in the lysosomal and inflammatory pathways.

A. Lysosomal pathway

B. Inflammatory pathway

Only Bonferroni's correct p values of <0.05 are shown

### Online Table 1

### Oligonucleotide Primers, Antibodies, and TaqMan probes

**Genotyping primers:**

| Gene | Forward sequence | Reverse sequence |
|---|---|---|
| *Cre recombinase* | TCTATTGCACACAGCAATCCA | CCAGCATTGTGAGAACAAGG |
| *Ctnnb1(Ex3)* | AGACTGCCTTGGGAAAAGCGC | TGGATGGGATCTGCATGCCCTCATCTA |
| *Ctnnb1(Ex8-13)* | TGCAGCTTACCTGACTCCTG | CCTGTTAGCCCTCATGGTGT |
| *Dsp* | GTTGGGCCTCTCGAATCAT | TCTTTGTCTGTTGCCATGTGA |

**List of antibodies**

| Antibodies | Concentration | Supplier | Catalog number |
|---|---|---|---|
| Anti-mouse IgG HRP linked antibody | 1:4000 (IB) | Cell Signaling Technology | 7076 |
| Anti-Rabbit IgG HRP linked antibody | 1:2000 (IB) | Cell Signaling Technology | 7074 |
| Anti-goat IgG HRP linked antibody | 1:2000 (IB) | R&D Systems | HAF109 |
| Donkey anti Rat Alexa Fluor 594 | 1:1000 (IF) | Invitrogen | A11007 |
| Goat anti Mouse Alexa Fluor594 | 1:1000 (IF) | Invitrogen | A11005 |
| Goat anti Rabbit Alexa Fluor 488 | 1:1000 (IF) | Invitrogen | A11012 |
| Vinculin | 1:10000 (IB) | Abcam | Ab129002 |
| CTNNB1 | 1:1000 (IB)<br>1:100 (IF) | Cell Signaling Technology | 8480S |
| TCF7L2 | 1:500 (IB) | Cell Signaling Technology | 2569S |
| TCF7L2 | 1:200 (IF) | Cell Signaling Technology | 2569S |
| DSP | 1:1000 (IB) | Abcam | ab71690 |
| DSP | 1:250 (IF) | Progen | 65146 |
| JUP | 1:1000 (IB) (IF) | Santa Cruz | 1497-R |
| PKP2 | 1:1000 (IB) | Santa Cruz | 18977 |
| PKP2 | 1:300 (IF) | Progen | 651101 |
| DSG2 | 1:500 (IB)<br>1:50 (IF) | Progen | 61002 |
| DSC2 | 1:200 (IB) | Santa Cruz | 70994 |
| DSC2 | 1:100 (IF) | Santa Cruz | 66863 |
| CASP3 | 1:1000 (IB) | Cell Signaling Technology | 14220 |
| RIPK1 | 1:1000 (IB) | Cell Signaling Technology | 3493 |
| RIPK3 | 1:1000 (IB) | Cell Signaling Technology | 15828 |
| MLKL | 1:1000 (IB) | Cell Signaling Technology | 37705 |

| | | | |
|---|---|---|---|
| GSDMD | 1:1000 (IB) | Cell Signaling Technology | 39754 |
| ASC | 1:1000 (IB) | Cell Signaling Technology | 67824 |
| COL1A1 | 1:300 (IF) | Abcam | 88147 |
| TGFB | 1:500 (IB) | R&D Systems | MAB1835 |
| SFRP2 | 1:100 (IHC) | R&D Systems | MAB1169 |
| SFRP4 | 1:200 (IHC) | Abcam | ab154167 |
| PCM1 | 1:1000 (IF) | Sigma | HPA023370 |
| PCM1 | 1:50 (IF) | Santa Cruz | Sc-398365 |
| AXIN2 | 1:1000 (IB) | ThermoFisher | pa5-25331 |
| cMYC | 1:1000 (IB) | Cell Signaling Technology | 5605S |
| DKK2 | 1:500 (IB) | R&D Systems | AF2435 |
| APOE | 1:1000 (IB) | Abcam | ab1906 |
| LGALS3 | 1:1000 (IB) | Abcam | ab53082 |
| Total OXPHOS Antibody Cocktail | 1:1000 (IB) | Abcam | Ab110413 |

**Oligonucleotide primers used in qPCR reactions**

| **Name** | **Sequence** |
|---|---|
| *Cox7a1* | Forward: TGCCAGGATCCGGAGTCTTAGAA<br>Reverse: GGTCATTGTCGGCCTGGAAG |
| *Atp5e* | Forward: TGGACTCAGCTACATCCGGT<br>Reverse: CGTTCGCTTTGAACTCGGTC |
| *Cox6a2* | Forward: GCCCAGAGTTCATCCCGTAT<br>Reverse: AGGATTGACGTGGGGATTGT |
| *Atp5g1* | Forward: CGCTCAGACCAAGGGCTAAA<br>Reverse: CCCTGGTACAGGAGCGAATC |
| *Cox7a2* | Forward: AGAGGACCATCAGCACCACT<br>Reverse: CATCAGATGCCCCGCCTTTC |
| *Cox8b* | Forward: GCCAGCCAAAACTCCCACTTC<br>Reverse: GCTCTCCAAGTGGGCTAAGA |
| *Cox6c* | Forward: GGCTGGTATCTTTCAGAGTGC<br>Reverse: TAGTTCAGGAGCGCAGGTCA |
| *Ndufa5* | Forward: AAGCTGGATATGGTCAAGGCG<br>Reverse: CTCTTCCACTTCACCACCCT |
| *Ndufs6* | Forward: TTTCGGGGTTCAAGTGTCGC<br>Reverse: CGAACCCTCCTGTAGTCTTT |
| *Uqcr10* | Forward: CGCAGAACTTCCACCTTTGC<br>Reverse: CCACAGTTTCCCCTCGTTGA |
| *Uqcrq* | Forward: GCCTTCCCAAGCTATTTCAGCA<br>Reverse: ACCTGGTTGCCCCATGTGTAG |
| *Vcl* | Forward: GGTCTAGCAAGGGCAATGAC<br>Reverse: TGAATAAGTGCCCGCTTGGT |
| *Lgals3* | Forward: TACTAGAAGCGGCCGAGC |

| | |
|---|---|
| | Reverse: TGTCTGCCATTTTCCTGGGTA |
| *Lrp1* | Forward: CAAAGCTGAAGGCTCCGAGT<br>Reverse: TATGCGGACACTCTCATCGC |
| *Tgfb1* | Forward: TGGAGCAACATGTGGAACTC<br>Reverse: GTCAGCAGCCGGTTACCA |
| *Tgfb2* | Forward: AGGAGTGGCTTCACCACAAAGACA<br>Reverse: ATTAGACGGCACGAAGGTACAGCA |
| *Tgfb3* | Forward: AGCTCTTCCAGATACTTCGACC<br>Reverse: AAAGACAGCCATTCAGCGGT |
| *Timp1* | Forward: CATGGAAAGCCTCTGTGGATA<br>Reverse: CTCAGAGTACGCCAGGGAAC |
| *Postn* | Forward: AGAGAAATCCCTGCACGACA<br>Reverse: GTTGGTGCAAACAAGGTCCA |
| *Col1a1* | Forward: TGACTGGAAGAGCGGAGAGTA<br>Reverse: GAGTAGGGAACACACAGGTCT |
| *Col3a1* | Forward: CTGTAACATGGAAACTGGGGAAA<br>Reverse: CCATAGCTGAACTGAAAACCACC |
| *Casp3* | Forward: AGCTTGGAACGGTACGCTAA<br>Reverse: GAGTCCACTGACTTGCTCCC |
| *Casp8* | Forward: GCGTGGAACAGGAAGTGAGTA<br>Reverse: GAAGAGCTGTAACCTGTGGC |
| *Bok* | Forward: CAGCGTATACCGGAACGTGG<br>Reverse: TTGCCCCATGTGATACCTGC |
| *Bak* | Forward: CAAGATCGCCTCCAGCCTATT<br>Reverse: CCCAGGAAGCCGGTCAAAC |
| *Bid* | Forward: CCGCAAACCTTTGCCTTAGC<br>Reverse: AACCGTTGCTGACCTCAGAGT |
| *Sfrp2* | Forward: CCCTTTGTAAAAATGACTTCGCAC<br>Reverse: CAGGATGATCTTGGTGTCTCTGT |
| *Spp1* | Forward: AGCAAGAAACTCTTCCAAGCAA<br>Reverse: GTGAGATTCGTCAGATTCATCCG |
| *Gdf15* | Forward: CTCAACGCCGACGAGTAC<br>Reverse: ACCCCAATCTCACCTCTGGA |
| *Mmp14* | Forward: GCCCTCTGTCCCAGATAAGC<br>Reverse: CCAGAACCATCGCTCCTTGA |
| *Mgp* | Forward: GCAACCCTGTGCTACGAATC<br>Reverse: CTTTTGGGCTTTAGCTCGCC |
| *Vim* | Forward: TGCACGATGAAGAGATCCAGG<br>Reverse: CTTTCATACTGCTGGCGCAC |
| *Mmp2* | Forward: TGCCCCCATGAAGCCTTG<br>Reverse: TGGTACAGCTGTTGTAGGAGG |
| *Ripk1* | Forward: TGACTTTCACATTAAGATAGCCGA<br>Reverse: AGTGGTGCTGCTCACTTCTTT |
| *Ripk3* | Forward: AGCTTTGGGATCCTCGTGTG<br>Reverse: TCTGTCAGTGGAGGACGACT |
| *Mlkl* | Forward: GTATTCAACAACCCCCAGGC<br>Reverse: CGCAAGATGTTGGGAGAATCG |
| *Gsdmd* | Forward: TTCCAGTGCCTCCATGAATGT<br>Reverse: ACAAACAGGTCATCCCCACG |
| *Asc* | Forward: GAGCAGCTGCAAACGACTAA |

| | |
|---|---|
| | Reverse: GACCCTGGCAATGAGTGCTT |
| *Casp1* | Forward: ACTGACTGGGACCCTCAAGT<br>Reverse: GCAAGACGTGTACGAGTGGT |
| *Zbp1 ex2-3* | Forward: GAGATGGGGCTCCTGCAATC<br>Reverse: CCCATTGGCTTCCAGGAATCTT |
| *Apoe* | Forward: CACACAAGAACTGACGGCAC<br>Reverse: CTCCAGCTCCTTTTTGTAAGCC |
| *Cox7b* | Forward: GCGATGTTGCCCTTAGCCA<br>Reverse: CTTGCCACCACTTGCTGAATG |
| *Ctsl* | Forward: ACCTTTAGTGCAGAGTGGCAC<br>Reverse: CCCCGTTGTGTAGCTGGAT |
| *Ctsd* | Forward: TACTCCATGCAGTCATCGCCT<br>Reverse: GACGACTGTGAAACACTGCG |
| *Ctsz* | Forward: GACCAGGCCGTTATCAACCA<br>Reverse: CCTTTCTCACCCCAGGGTTC |
| *Ctsa* | Forward: CACCTTGGCTGTACTGGTCAT<br>Reverse: AGTAGACCAGGGAGTTGTCGT |
| *Acp5* | Forward: GACCCACCGCCAAGATGGAT<br>Reverse: CACAAATCTCAGGGTGGGAGT |
| *Man2b1* | Forward: CCGTCGCTTCATCTATGTGGA<br>Reverse: ATCACCCAGCCACCATTGAC |
| *Slc17a5* | Forward: TCTTCACCCTGTTCACACCG<br>Reverse: AAGACCACATGGCGTGCATA |
| *Gm2a* | Forward: CTCACGATCCAACCTGACCC<br>Reverse: ATCTTGACCCAGAAGCCAGC |
| *Ccl2* | Forward:TTAAAAACCTGGATCGGAACCAA<br>Reverse: GCATTAGCTTCAGATTTACGGGT |
| *Cd44* | Forward:GGCTCTGATTCTTGCCGTCT<br>Reverse:TCCTGTCTTCCACTGTCCCA |
| *Tnc* | Forward:ATGGCTGTTGTTGCTATGGCA<br>Reverse: ACGGCTACCACAGAAGCTG |
| *Il6* | Forward: AGCCAGAGTCCTTCAGAGAGAT<br>Reverse: GAGAGCATTGGAAATTGGGGT |
| *Tlr5* | Forward: GACGCCTCATCTCACTGCAT<br>Reverse: ATGTTCCAAGCGTAGGTGCT |
| *Tlr6* | Forward: GGTACCGTCAGTGCTGGAAAT<br>Reverse: GGGCGCAAACAAAGTGGAAA |
| *Itgam* | Forward: GTGTGACTACAGCACAAGCC<br>Reverse: GGACAGGCCCAAGGACATATT |
| *Itgal* | Forward: TGAAGATGGGGTTGTCGTGG<br>Reverse: AACCATGTAGGCTGACTGGC |
| *H2-DMa* | Forward: TGGAGCATGAGCAGAAGTCAG<br>Reverse: CTGTGGGTCATCCCAGAACA |
| *H2-Eb1* | Forward:GCGGAGAGTTGAGCCTACG<br>Reverse: CCAGGAGGTTGTGGTGTTCC |
| *Cybb* | Forward: AAAACTCCTTGGGTCAGCAC<br>Reverse: AGATTTCGACACACTGGCA |
| *Dpep2* | Forward: ATGCAGGAGTTCCCGCTTAT<br>Reverse: GGTCTGGCCATGGGTGAAG |

| | |
|---|---|
| *IL6ra* | Forward: AATGATCACCTGGTGGGGAC<br>Reverse: CACTCACAGATGGCGTTGAC |
| *H2-DMb1* | Forward: GCCCTTCTGGAATGCGCTG<br>Reverse: CCGGCTCCCTTGTGTTAAAAG |
| *Hmox1* | Forward:AAGCCGAGAATGCTGAGTTCA<br>Reverse:GCCGTGTAGATATGGTACAAGGA |

**TaqMan probes**

| **Gene** | **TaqMan Assay ID** |
|---|---|
| *Vcl* | Mm00447745_m1 |
| *Myh7* | Mm0060555_m1 |
| *Nppb* | Mm01255770_g1 |
| *Nppa* | Mm01255748_g1 |
| *Acta1* | Mm00808218_g1 |
| *Atp2a2* | Mm00437634_m1 |
| *Myh6* | Mm00440354_m1 |
| *Ctgf* | Mm01192932_g1 |
| *Ctnnb1* | Mm00483039_m1 |

Online Table 2

Summary of STAR alignment of the RNA Sequence Reads

| Sample | Genotypes | Total number of reads | Uniquely mapped reads (N) | Uniquely mapped reads (%) |
|---|---|---|---|---|
| 1-6821-WT-S1-L001 | Wild type | 56,986,661 | 44,080,962 | 77.35 |
| 2-6840-WT-S2-L001 | Wild type | 58,592,357 | 45,042,018 | 76.87 |
| 3-6841-WT-S3-L001 | Wild type | 57,741,248 | 43,997,161 | 76.2 |
| 4-6875-WT-S4-L001 | Wild type | 47,657,324 | 36,515,508 | 76.62 |
| 5-6876-WT-S5-L001 | Wild type | 51,350,429 | 36,020,507 | 70.15 |
| 21-6670-DSP-S19-L003 | *Myh6-Mcm*$^{Tam}$:*Dsp*$^{F/F}$ | 47,588,510 | 36,116,114 | 75.89 |
| 22-6699-DSP-S26-L004 | *Myh6-Mcm*$^{Tam}$:*Dsp*$^{F/F}$ | 53,655,861 | 42,741,828 | 79.66 |
| 23-6701-DSP-S27-L004 | *Myh6-Mcm*:*Dsp*$^{F/F}$ | 48,350,386 | 36,325,713 | 75.13 |
| 24-6731-DSP-S28-L004 | *Myh6-Mcm*$^{Tam}$:*Dsp*$^{F/F}$ | 61,200,040 | 50,444,975 | 82.43 |
| 25-6818-DSP-S22-L004 | *Myh6-Mcm*$^{Tam}$:*Dsp*$^{F/F}$ | 59,729,286 | 47,836,233 | 80.09 |
| 31-6679-LOFDSP-S34-L005 | *Myh6-Mcm*$^{Tam}$:*Dsp*$^{F/F}$:*Ctnnb1*$^{LoF}$ | 54,350,416 | 41,001,508 | 75.44 |
| 32-6706-LOFDSP-S35-L005 | *Myh6-Mcm*$^{Tam}$:*Dsp*$^{F/F}$:*Ctnnb1*$^{LoF}$ | 47,519,932 | 37,832,555 | 79.61 |
| 33-6742-LOFDSP-S29-L005 | *Myh6-Mcm*$^{Tam}$:*Dsp*$^{F/F}$:*Ctnnb1*$^{LoF}$ | 60,049,335 | 47,080,290 | 78.4 |
| 34-6754-LOFDSP-S30-L005 | *Myh6-Mcm*$^{Tam}$:*Dsp*$^{F/F}$:*Ctnnb1*$^{LoF}$ | 50,392,020 | 38,836,949 | 77.07 |
| 35-6887-LOFDSP-S31-L005 | *Myh6-Mcm*$^{Tam}$:*Dsp*$^{F/F}$:*Ctnnb1*$^{LoF}$ | 61,055,959 | 47,533,308 | 77.85 |
| 26-6702-GOFDSP-S23-L004 | *Myh6-Mcm*$^{Tam}$:*Dsp*$^{F/F}$:*Ctnnb1*$^{GoF}$ | 53,099,088 | 41,495,349 | 78.15 |
| 27-6799-GOFDSP-S24-L004 | *Myh6-Mcm*$^{Tam}$:*Dsp*$^{F/F}$:*Ctnnb1*$^{GoF}$ | 52,853,929 | 41,722,460 | 78.94 |
| 28-6808-GOFDSP-S25-L004 | *Myh6-Mcm*$^{Tam}$:*Dsp*$^{F/F}$:*Ctnnb1*$^{GoF}$ | 55,272,569 | 42,842,348 | 77.51 |
| 29-6896-GOFDSP-S32-L005 | *Myh6-Mcm*$^{Tam}$:*Dsp*$^{F/F}$:*Ctnnb1*$^{GoF}$ | 53,221,699 | 42,538,183 | 79.93 |
| 30-6899-GOFDSP-S33-L005 | *Myh6-Mcm*$^{Tam}$:*Dsp*$^{F/F}$:*Ctnnb1*$^{GoF}$ | 58,226,579 | 44,829,357 | 76.99 |

Data in the wild type and *Myh6-Mcm*$^{Tam}$:*Dsp*$^{F/F}$ groups have been published. (1)

1. Olcum MR, L.; Fan, S.; Gonzales, M.M.; Jeong, H.H.; Zhao, Z.; Gurha, P.; Marian, A.J. PANoptosis is a prominent feature of desmoplakin cardiomyopathy. *J Cardiovasc Aging.* 2023;3(3):1-20.
